# Genome editing in mammals using CRISPR type I-D nuclease

**DOI:** 10.1101/2020.03.14.991976

**Authors:** Keishi Osakabe, Naoki Wada, Emi Murakami, Yuriko Osakabe

## Abstract

Adoption of the CRISPR-Cas system has revolutionized genome engineering in recent years; however, application of genome editing with CRISPR type I—the most abundant CRISPR system in bacteria—has been less developed. Type I systems in which Cas3 nuclease degrades the target DNA are known; in contrast, for the sub-type CRISPR type I-D (TiD), which lacks a typical Cas3 nuclease in its cascade, the mechanism of target DNA degradation remains unknown. Here, we found that Cas10d—a nuclease in TiD—is multi-functional in PAM recognition, stabilization and target DNA degradation. TiD can be used for targeted mutagenesis of genomic DNA in human cells, directing both bi-directional long-range deletions and short insertions/deletions. TiD off-target effects, which were dependent on the mismatch position in the protospacer of TiD, were also identified. Our findings suggest TiD as a unique effector pathway in CRISPR that can be repurposed for genome engineering in eukaryotic cells.

## INTRODUCTION

CRISPR-Cas (clustered regularly interspaced short palindromic repeats-CRISPR-associated) systems are RNA-based adaptive immune systems that protect bacteria and archaea from viruses, plasmids, and other foreign genetic elements. The CRISPR systems, especially, Class 2 type II systems [CRISPR/Cas9 (Cong et al, 2013; Mali et al, 2013)] and type V [CRISPR/CpfI (Zetsche et al. 2015)] have been widely used to efficiently introduce a mutation on a target DNA of interest in eukaryotic chromosomal DNA as a tool of genome editing. Although less common than Class 2 systems, Class 1 type I CRISPR systems have some advantages compared to Cas9 and Cpf1 as genome editing tools, such as a large variety of mutation profiles, including longer gRNA sequences, which might allow higher specificity and long-range genome deletion (Cameron et al. 2019; Dolan et al. 2019; Morisaka et al. 2019).

Class 1 type I CRISPR systems include a wide variety of subtypes. Type I systems comprise a guidance short RNA molecule, the so-called CRISPRs RNA (crRNA), a target recognition module, and a DNA cleavage module. In nature, crRNAs are transcribed from the CRISPR array in precursor form, and are subsequently processed into their mature forms, typically about 30–50 nucleotides in length (Wang et al. 2011; Shao and Li. 2013; Niewoehne et al. 2014). After processing of crRNA by Cas6, a mature crRNA is incorporated into the target recognition module, which consists of Cas5, Cas6, Cas7, and Cas8, the so-called Cascade (CRISPR-associated complex for antiviral defense) (Gleditzsch et al. 2016; Hayes et al. 2016; Hochstrasser et al. 2016). Cascade complexes recognize short (typically 3–5 base) sequence elements called protospacer adjacent motifs (PAMs) in foreign DNA (Mojica et al. 2009; Shah et al. 2013; Leenay et al. 2016). Components of Cascade differ in type I sub-families, varying from three Cas effectors (Cas5, Cas7 and Cas8) in type I-C (Hochstrasser et al. 2016) to five Cas effectors (Cas5, Cas6, Cas7, Cas8, and Cse2) in type I-E (Hayes et al. 2016). In type I systems, the Cascasde complexe disrupt local DNA pairing to form R-loop structures, in which base-pairs between crRNA and the complementary target strand, displacing the non-target DNA strand (Sashital et al. 2012; Anders et al. 2014; Hochstrasser et al. 2014; Hayes et al. 2016; Xiao et al. 2018). This binding and unwinding of the double-stranded DNA by the Cas effectors and crRNA is required for DNA cleavage and destruction by type-specific Cas effector nucleases Cas3 (Sapranauskas et al. 2011; Sinkunas et al. 2011, 2015; Xiao et al. 2018). Cas3 and Cas3’ are known as DNA cleavage modules in the type I system. Both proteins possess a central helicase motif, and, most importantly, Cas3 also has a nuclease domain, the so-called HD domain. Recent studies have shown that Cas3 is recruited to the R-loop formed by Cascade, and that it nicks non-target strand DNAs, processively degrading DNA in the region upstream of the PAM-proximal side (Huo et al. 2014; Xiao et al. 2018; Dolan et al. 2019).

We identified a CRISPR genomic locus encoding a previously assigned, but uncharacterized, Class 1 type I subtype: type I-D (TiD). CRISPR TiD contains a Cas3 effector protein, but lacks a nuclease domain; instead, Cas10d—a unique effector protein of TiD not found in other CRISPR systems— possesses a typical nuclease domain. In addition, one of major Cascade factors for PAM recognition, Cas8 homologous protein (Cass et al. 2015) is missing in TiD. Therefore, the mode of interference through PAM recognition to target DNA degradation in the TiD system will differ from that of other type I systems. Here, we explore the function of novel Cas effector proteins from TiD, identifying Cas10d protein as the functional nuclease in this system. TiD crRNA was determined as a 35–36 base target recognition sequence that binds to five Cas proteins, yielding a complex that is critical for crRNA function. Of particular importance, we found that Cas10d acts both as the functional nuclease effector protein and in PAM recognition through interaction with the TiD cascade, including Cas3d.

Furthermore, engineered TiD crRNA generated mutations in target DNA in human cells introduced small insertions/deletions (indels) but also long-range and bi-directional DNA deletions in the target genome; TiD-induced mutation patterns differ from those of other CRISPR systems. We also analyzed a unique feature of off-target mutation against a mismatch position in the TiD gRNA. Our findings suggest novel genome editing tools for eukaryotic cells using the CRISPR effector module pathway of the TiD system.

## RESULTS AND DISCUSSION

### TiD is composed of Cas effector proteins with a nuclease module

Components of Cas effector proteins and the crRNA sequence from *Microcystis aeruginosa* were evaluated using BLAST. The 7.6 kb CRISPR/Cas TiD locus of *M. aeruginosa* strain PCC9808 consists of eight Cas genes (Cas1d–Cas7d, Cas10d) followed by an array of 36 repeat-spacer units (**Fig. 1A**). The HD domain that functions in DNA cleavage in CRISPR type I-A, B, C, E, and F (Sinkunas et al. 2015; Hochstrasser et al. 2016; Li et al. 2016; Pyne et al. 2016; Guo et al. 2017) is lacking in Cas3d (Gong et al. 2014); however, Cas10d harbors an HD-like nuclease domain in its N-terminal region (**Fig. 1B, Supplemental Fig. S1**). Although Cas10 proteins in CRISPR type III possess the HD domain (Mogila et al. 2019), Cas10d in TiD was highly divergent to Cas10s in type III, rather the HD domain of TiD was similar to the HD domain of Cas3 proteins in CRISPR type I-B, C, E, and F **(Supplementary Fig. S1).** To confirm whether the Cas10d has nuclease function, we performed *in vitro* nuclease assays. *In vitro*-synthesized Cas10d protein was able to cleave M13 ssDNA as a substrate in the presence of ATP and Ni^2+^ and Co^2+^ (**Fig. 1C**), suggesting that it acts as a nuclease in the TiD system. The typical Cas8 gene— the common effector in CRISPR type I-A, B, C, E, and F (Cass et al. 2015; Makarova et al. 2015)—is missing from CRISPR/Cas TiD locus of *M. aeruginosa*, predicting different mechanisms of cascade complex stability and *in vivo* DNA cleavage activity of TiD compared to other type I sub-types. To identify the PAM on the TiD system in *M. aeruginosa*, we performed a depletion assay using the negative selection marker *ccdB*. pCmMa567Δ10 (**Supplemental Fig. S6**), containing expression cassettes for Cas5d, Cas6d, Cas7d and Cas10d carrying a mutation in the HD-like domain [Cas10d(H177A)] and gRNAs targeted to the *ccdB* promoter, was introduced into *Escherichia coli* strain BL21-AI (**Fig. 1D**) followed by a PAM library plasmid, pPAMlib-ccdB (**Supplemental Fig. S6**). *ccdB* negative selection revealed the PAM: 5’-GTH-3’ (H=A or C or T) adjacent to the target sequence (**Fig. 1D, right panel**). When pCmMa567, which carries expression cassettes for Cas5d, Cas6d and Cas7d, was used for screening instead of pCmMa567Δ10, GTH PAMs were screened out, but the resulting transformants were unstable, and growth of the *E. coli* cells was very weak. These results suggested that Cas10d requires the correct PAM for full repression, and that Cas10d is a functional counterpart of Cas8 for PAM recognition and stabilization. We did not find any similar amino acid sequences shared between the Cas10d and Cas8 protein families. The precise structural features of Cas10d should be determined to shed further light on the key role of Cas10d.

**Figure 1.**
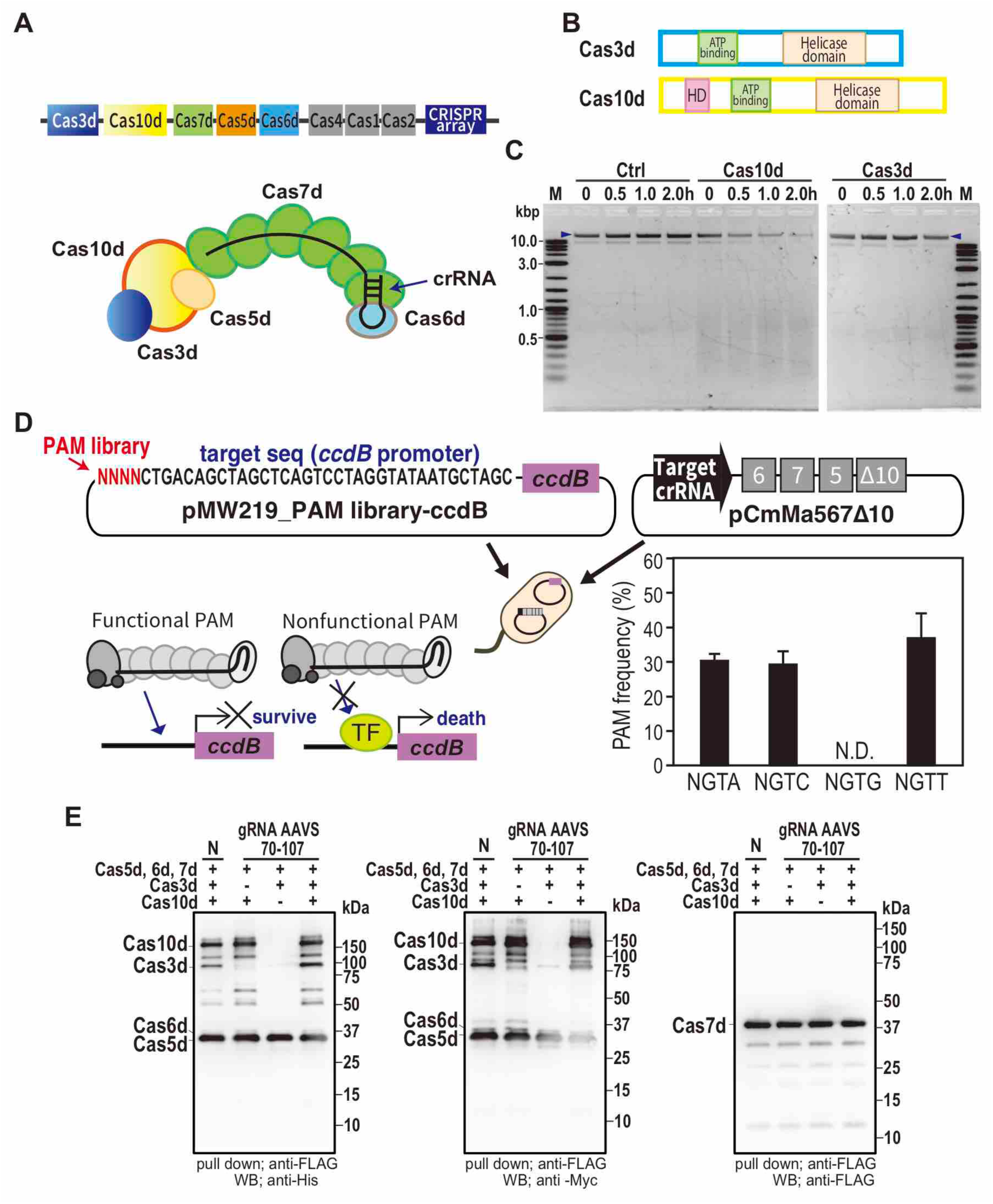
DNA cleavage activity of CRISPR type I-D. **(A)** The CRISPR type I-D (TiD) structure. Upper; the CRISPR type I-D locus in *M. aerginosa*. Lower; subunit organization of TiD and schematic of crRNA (black) of TiD. **(B)** The HD nuclease domain of Cas10d. **(C)** The detection of Cas10d activity with in-vitro DNA cleavage assay. M13mp18 single-strand DNA (NEB) was used for the cleavage assay. Ctrl; control reaction without Cas proteins, blue arrows; undigested DNA. **(D)** PAM identification by the *E. coli* negative selection screening using *ccdB* expression system. PAM library was inserted in front of the target sequence of *ccdB* promoter. PAM frequency was determined using the survived *E. colli* cells. Data are means ± S.E. of independent experiments (n = 3). **(E)** in vitro pull-down assay of TiD Cas proteins containing the target gRNA sequence (*AAVS* 70-107, **Supplemental Table S1**). Each Cas expression vector (Flag-NLS-Cas7d, Myc-bpNLS-Cas5d-bpNLS-6xHis, Myc-bpNLS-Cas6d-bpNLS-6xHis, Myc-bpNLS-Cas7d-bpNLS-6xHis, Myc-bpNLS-Cas10d-bpNLS-6xHis) and gRNA were infected into HEK293T cells and the pull-down assay using the cell lysates was performed with the antibodies as described in the figures.

### TiD is biologically active in human cells as a genome editing tool

To confirm protein expression of heterogenous host cells and complex formation of TiD Cas proteins, Cas genes were expressed in HEK293T cells, and *in vivo* complex formation of Cas3d, Cas5d, Cas6d, Cas7d, and Cas10d with crRNA was shown by pull-down assay (**Fig. 1E**). Importantly, Cas3d could not bind to the TiD Cascade without Cas10d.

Genome editing activity of the TiD complex was evaluated using the luciferase single-strand annealing (SSA) recombination system consisting of NanoLuc luciferase (Hall et al. 2012) containing a 300 bp homologous arm separated by a stop codon and the human *AAVS1* and *EMX1* gene fragments containing the TiD target site (gRNA target sequences listed in **Supplemental Table S2**). HEK293T cells transfected simultaneously with TiD Cas gene and TiD crRNA expression vectors were screened by luc reporter assay 72 hr after transfection (**Fig. 2A**). Expression of TiD led to notably higher (2- to 4-fold) activity than the non-target control (**Fig. 2A-D**). Wild-type Cas10d and Cas10d(H177A) were then tested in the luciferase reporter assay using gRNAs *AAVS* GTC_70–107(+) and *AAVS* GTC_159– 196(+) (**Fig. 2B, Supplemental Table S1**). The results indicated that Cas10d indeed possesses a nuclease activity that can be utilized for genome editing in human cells. In the original CRISPR locus of *M. aeruginosa* strain PCC9808, both 35 b and 36 b protospacer sequences are used to target specific genomic DNAs; both gRNAs functioned for genome editing in human cells (**Fig. 2C**). TiD activity also required the expression of pre-mature crRNA with gRNA (**Fig. 2D**). Cas6 is known as a endoribonuclease, and cleaves repeat sequences of the pre-mature crRNA at specific positions and hold the repeat-derived 3’ handle of the mature crRNA (Wang et al. 2011; Shao and Li. 2013; Niewoehner et al. 2014; Jesser et al. 2019). Processing of pre-crRNA into its mature form might be important for stabilizing the crRNA interaction with Cascade and for functional activity in vivo by TiD. We also evaluated the effects of the position and number of nuclear localization signals (NLS) by using two types of NLS: monopartite-NLS (SV40NLS) and bipartite-NLS (bpNLS). bpNLS functioned as effectively on both N-ter and C-ter in Cas3d, Cas5d, Cas6d, and Cas10d as SV40NLS did on N-ter (**Supplemental Fig. S3A, B)**; however, bpNLS attached to Cas7d disrupted TiD activity, although it was expressed strongly in the nucleus (**Supplemental Fig. S3C**).

**Figure 2.**
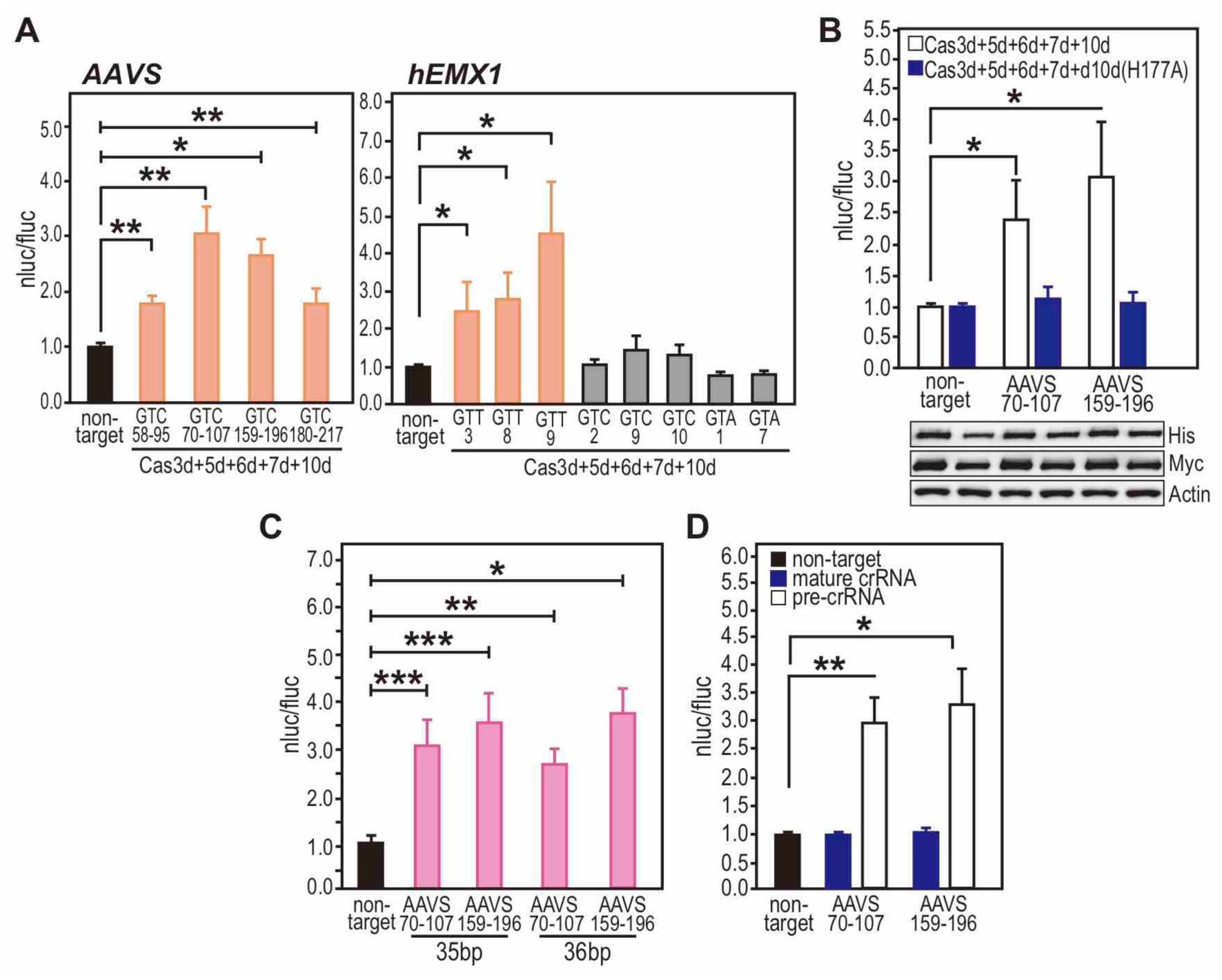
Detection of CRISPR TiD-mediated genome editing by LUC reporter assay. **(A)** The Cas expression vectors and a LUC reporter vector harboring the target sequence were transfected into HEK293T cells; an endonuclease cleavage assay produced luminescence. The charts show selection of gRNAs for *AAVS* and *hEMX1* genes in the TiD system using the luc reporter assay. Data are means ± S.E. of independent experiments (n = 4). *P < 0.05 is determined by Student’s *t* tests. **(B)** Effect on the HD domain of Cas10d. The luc reporter assay was performed using the *AAVS* GTC_70-107(+), *AAVS* GTC_159-196(+), and non-target gRNAs. Upper; White bars; luc reporter assay of the TiD system with the wild-type Cas10d, black bars; luc reporter assay of the TiD system with the HD-mutated dCas10d (H177A). Data are means ± S.E. of independent experiments (n = 4). Asterisks indicate statistically significant differences determined by Student’s t tests. *P < 0.07. Lower; The expression levels of dCas10 and dCas10d detected by the antibodies for the tags fused to the N- (Myc) or C-ter (His). For the Cas10d and dCas10d, pEFs-Myc-bpNLS-dCas10 or dCas10d (H177A)-bpNLS-6xHis and pEFs-SV40NLS HA-Cas3d, Strept-Cas5d, Myc-Cas6d, or FLAG-Cas7d was used for the luc reporter assay. **(C)** Effect of the gRNA target sequence length in the TiD activity. Data are means ± S.E. of independent experiments (n = 4). *P < 0.005, **P < 0.01, and ***P < 0.05 are determined by Student’s t tests. **(D)** Effect of the crRNA structure used in the TiD system. Data are means ± S.E. of independent experiments (n = 4). *P < 0.01 and **P < 0.05 are determined by Student’s t tests.

### Impact on TiD activity of mismatch sequence between protospacer and target DNA

We explored the mismatch effect of spacer sequence on genome editing activity using the luc reporter assay (**Fig. 3, Supplemental Fig. S2**). The 23 nucleotides just adjacent to the PAM, with the exception of nucleotides at 1-, 6-, 12-, and 18-nt downstream of PAM, were identified as important for genome editing activity; however, a single mismatch at 24–36 nt showed no effect on genome editing activity (**Fig. 3A, B upper panel**). Previous studies have shown that the 6th nucleotide of the protospacer is not engaged in base pairing with the target DNA strand, because every 6th nucleotide binds with the thumb domain of Cas7 subunits in the Cascade complex (Jackson et al. 2014; Mulepati et al. 2014; Zhao et al. 2014; Xiao et al. 2018). The obvious difference between TiD and type I-E was the effect of a mismatch at 1-nt, where the DNA cleavage activity in the type I-E system is markedly reduced (Morisaka et al. 2019), suggesting that the mode and/or position of RNA-Cas7 binding might differ between the two systems. In addition, no homolog of small subunit Cse2 in type I-E was found in the TiD locus. In the type I-E system, Cse1 and two subunits of Cse2 function to stabilize the entire R-loop (**Fig. 3C**). This structural difference in Cascade between type I-D and type I-E might affect the mismatch effect of each system.

**Figure 3.**
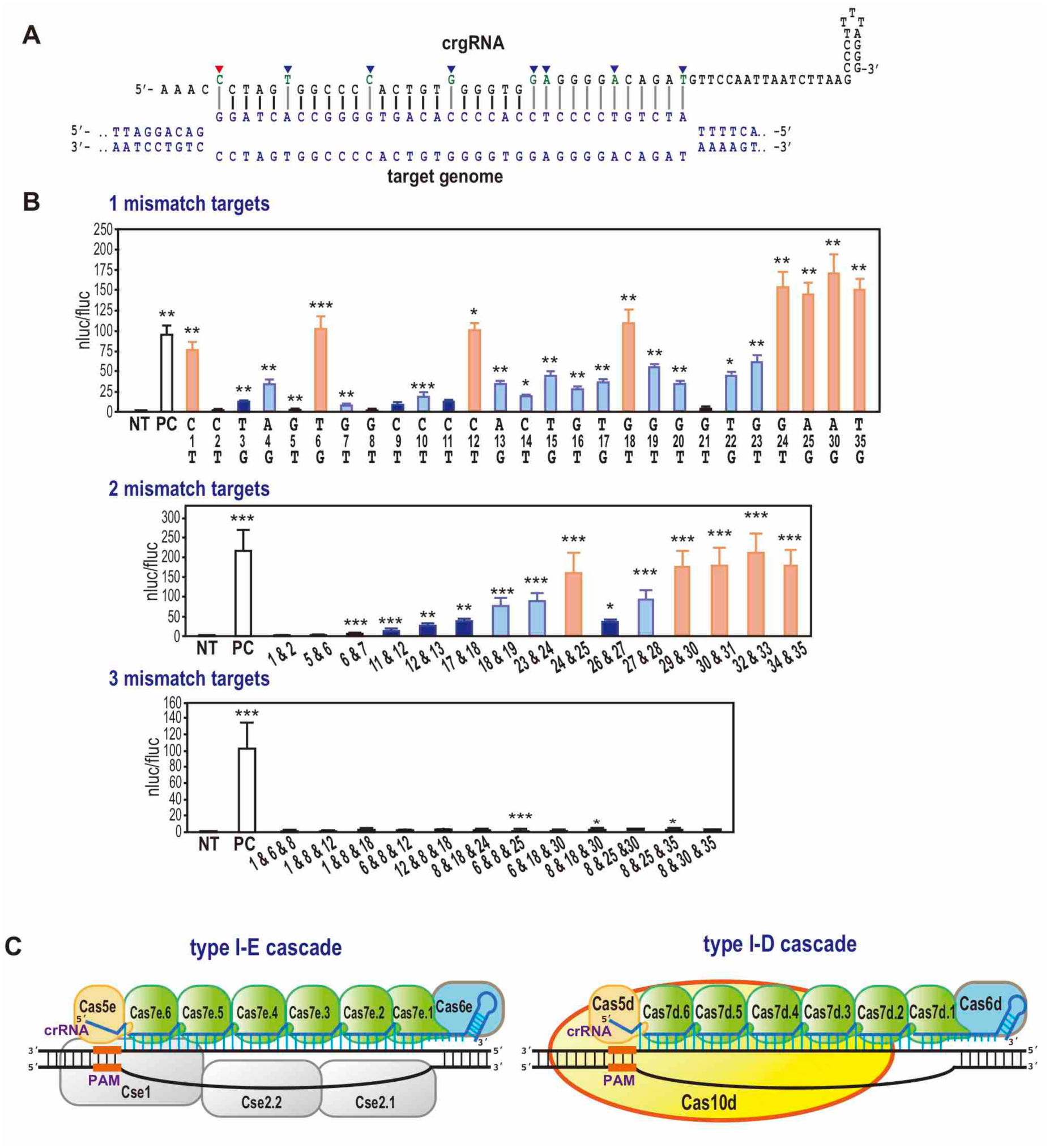
Detection of off target effects of CRISPR TiD mediates genome editing. **(A)** Evaluation of ozzff-target effects in the TiD gRNA. The critical nucleotides in the target gRNA sequence of *AAVS* GTC_70-107(+) for genome editing activity of TiD were evaluated using the luc reporter assay in human HEK293T cells, which was performed using the split-type vectors (**Supplemental Methods**). The nucleotides indicated with blue arrows; the nucleotides at 6-, 12-, 18-, and 24-nt positions that weakly affect the TiD activity. The first nucleotide (red arrow) just adjacent PAM also has a weak effect to the TiD activity. NT; non-target, PC; the positive control with the non-mutated gRNA. Data are means ± S.E. of independent experiments (n = 3). *P < 0.001, **P < 0.005, and ***P < 0.01 are determined by Student’s t tests and statistically significant differences compared with NT. **(B)** Nucleotides in the target gRNA sequence of *AAVS* GTC_70-107(+) containing two or three mismatches that are critical for genome editing activity of TiD were evaluated using the luc reporter assay in human HEK293T cells using split-type vectors. Numbers indicate the mismatch positions in the gRNA target sequences. NT; non-target, PC; positive control with non-mutated gRNA. Data are means ± S.E. of independent experiments (*n* = 3). *P < 0.005, **P < 0.01, and ***P < 0.05 were determined by Student’s *t* tests and statistically significant differences compared with NT. **(C)** Comparison of type I-D, E, and F cascade and R-loop formation. Mechanisms of target DNA recognition and components of type I-E and F have been deduced from structural analyses (Pausch et al. 2017). CRIPSR Type I-D does not have Cse1 and Cse2; instead, Cas10d could hold the R-loop to digest the target sequences.

We further analyzed the effect of mismatch number and position in the gRNA sequence.

Introduction of two mismatches within the region covered by nucleotides 1 to 28 markedly reduced genome editing activity except for the combination of mismatches at position 24-nt and 25-nt; however, within nucleotide positions 29-nt to 35-nt, two mismatches had less effect of reduction of TiD activity (Fig. 3B middle panel). On the other hand, the introduction of three mismatches completely abolished TiD activity (Fig 3 lower panel). These findings highlight important aspects of the specific interaction between protospacer and target DNAs, and the subsequent DNA cleavage mechanisms of TiD; however, further studies, including structural analysis of TiD, will be needed to understand the finer details of the system.

### Targeted-mutagenesis by TiD in HEK293 cells

We next targeted an internal genomic DNA region corresponding to the TiD gRNA target. We constructed single expression vectors for each Cas gene, a vector containing a triple-gene expression cassette connected via a 2A self-cleavage peptide sequence (2A), and a vector containing a quintuple gene expression cassette connected via 2A. Of all these configurations, transfection with the single Cas gene expression vectors yielded the highest Cas protein expression levels in human cells (**Supplemental Fig. S4**).

Following a recent genome editing study using CRISPR/Cas type I-E (Dolan et al. 2019), we speculated that long-range deletion might be introduced near the TiD target site. To investigate this, we selected targets *hEMX1* GTT9(–) for the human *EMX1* gene and *AAVS* GTC_70–107(+) for the *AAVS* gene based on the luciferase SSA assay (**Fig. 2A**). Using two primers flanking 5–19 kb around the and *AAVS* and *hEMX1* target sites (**Fig. 4A, B**), DNA fragments were PCR-amplified from total DNA from HEK293T cells harboring the TiD system, and cloned. Sequencing revealed that TiD introduced long-range deletions ranging from 5 kb to over 19 kb at both target sites (**Fig. 4A, B, C**). *AAVS* GTC_70– 107(+) mutated sequences shared some specific features: the size of major long-range deletions was not random but showed some restrictions in the case of the target *AAVS* GTC_70–107(+) as 5.2–5.5 kb and 17 kb deletions; with some minor exceptions, mostly bi-directional deletions were detected (**Fig. 4A, B, C, Supplemental Fig. S5**). These features differed from those following mutation by type I-E (Dolan et al. 2019), and type II effectors such as Cas9 (Cong et al, 2013; Mali et al, 2013) and Cpf1 (Zetsche et al. 2015). Microhomology and insertions were also observed in TiD mutation sites (**Fig. 4A, Supplemental Fig. S5**). The mutation rates for long-range deletion by TiD in the cloned DNA fragments were 55.0% and 57.1% with *hEMX1* GTT_9(–) and *AAVS* GTC_70–107(+), respectively. Furthermore, the mutation distribution in cloned PCR products varied among targets; mutations by *hEMX1* GTT_9(–) showed low mosaic, whereas mutations by *AAVS* GTC_70–107(+) showed more varied mutated sequences (**Fig 4A, B, Supplemental Fig. S5**).

**Figure 4.**
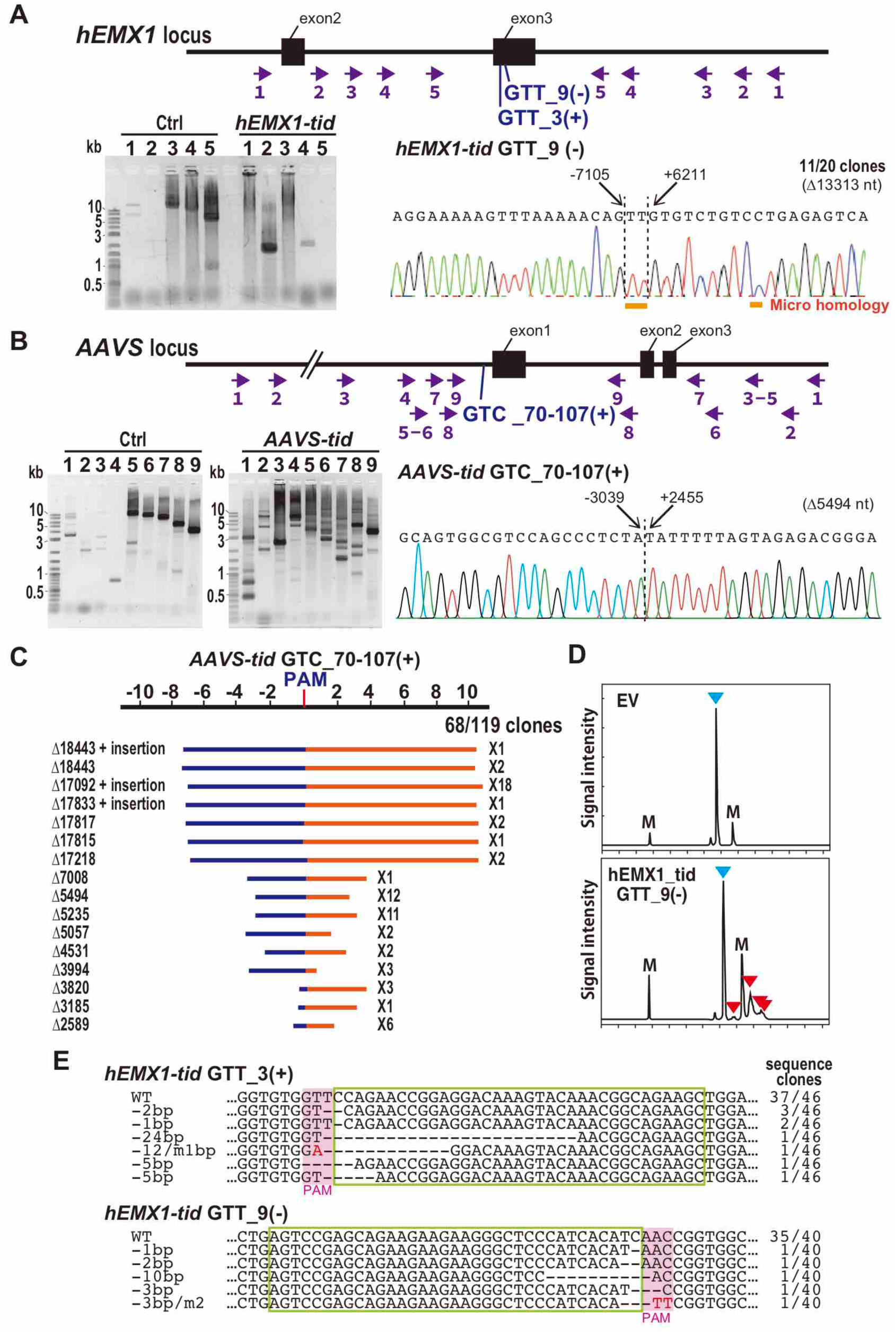
CRISPR TiD mediates genome editing in human cells. **(A) (B)** The detection of long-range deletion mutations in the *EMX1* and *AAVS* genes induced by the CRISPR TiD. Upper; gene structure, gRNA positions (arrows), and the various primer sets to amplify the mutation. Lower (left); The PCR amplified fragments separated on agarose gels. Numbers show the primer sets shown in the gene structures. Lower (right); The large deletion mutations analyzed by the Sanger sequencing of the cloned DNA from the CRISPR TiD infected cells. The nucleotide positions from the PAM were indicated on the sequence. **(C)** The long-range deletion pattern induced by *AAVS* GTC_70-107(+). Blue; deletion in the 5’ upstream from the PAM, red; deletion in the 3’ downstream from the PAM. The distribution of deletion mutation was also shown in **Supplemental Fig. S5A.** **(D)** Heteroduplex mobility assay of the mutation in the *hEMX1* gene induced by CRISPR TiD. Multiple heteroduplex peaks (red arrows) were detected in PCR amplicons from the HEK293T cells infected with CRISPR TiD that targets to *hEMX1* gene (GTT_9). blue arrow; Wild-type peak. EV; the cells infected with empty vector. M; internal markers. **(E)** Mutation sequences in HEK293T transfected with CRISPR TiD that target to *hEMX* gene. WT; wild-type sequences. Two gRNAs for CRISPR TiD (GTT_3 and GTT_9) were separately infected to HEK293T cells. gRNA target sequences are indicated in green boxes and PAM is indicated in pink boxes. The sequence frequencies in the cloned PCR products were indicated in the right of the sequence.

The luciferase SSA assay, with a short (300–500 bp) fragment spanning two homologous regions of the luciferase gene, clearly showed the genome editing activity of TiD, thus we next asked whether, like Cas9 and Cpf1, TiD could also introduce small changes at the target site. To test this, the 300–500 bp target region was PCR-amplified and analyzed by heteroduplex mobility assay (HMA). Typical multiple heteroduplex HMA peaks were detected in PCR amplicons from HEK293FT cells transfected with TiD with *hEMX1* GTT_9(–)gRNA (**Fig 4D**). Sequence analysis of cloned PCR products from HEK293FT cells transfected with gRNAs *hEMX1* GTT_3(+) and *hEMX1* GTT_9(–) showed that gRNA coupled with TiD Cas proteins could introduce double-strand breaks (DSBs) and small indels at the target site at mutation rates of 19.6 % [*hEMX1* GTT_3(+)] and 12.5% [*hEMX1* GTT_9(–)] in the cloned DNAs (**Fig. 4E**).

## Conclusions

Detailed studies on prokaryote genomes have identified eight subtypes of CRISPR type I families to date (Makarova et al. 2015), but other subtypes, such as CRISPR TiD, remain less well characterized. A unique member of the type I subtype, CRISPR TiD does not possess a Cas3 nuclease but instead has a Cas10d nuclease, which is the functional component of TiD. In this study, we developed the first CRISPR TiD system for use as a genome editing tool, and showed efficient site-directed mutagenesis in human cells. Notably, in the TiD system, Cas10d—a unique Cas effector protein of TiD—is the multifunctional effector for PAM recognition, stabilization and target DNA degradation. The DNA cleavage mechanism of TiD and the subsequent DNA repair pathway may differ from those of Cas3, Cas9, and Cpf1. The ability of type I CRISPR to generate such a diverse range of large deletions from a single targeted site could potentially enable long-range chromosome engineering that would allow simple and effective multi-gene function screening. As a novel technology in the CRISPR toolbox, TiD opens new possibilities in genome engineering.

## MATERIALS AND METHODS

### Vector construction

Details of all plasmid DNAs used in this study are described in **Supplemental Fig. S6.** PCR amplification for cloning gene fragments was done using PrimeSTAR Max (TaKaRa). Cloning for assembling was done using Quick ligation kit (NEB), NEBuilder HiFi DNA Assembly (NEB), and Multisite gateway Pro (Thermo Fisher Scientific).

#### Bacterial vectors

Gene fragments corresponding to *E. coli* codon-optimized *Cas* effector genes consisted of Cascade; *Cas5d, Cas6d, Cas7d* and the expression cassette for crRNA were synthesized (gBlocks®) (IDT), assembled, and cloned into the pACYC184 vector (Nippon Gene) separately. Expression of *Cas* genes and crRNAs was driven by the T7 promoter. For crRNA expression, a DNA fragment containing a T7 promoter-repeat-spacer-repeat sequence was cloned into pACYC184. After confirming sequences of each gene expression cassette, *Cas* gene and crRNA cassettes were re-assembled into pACYC184 to yield pCmMaTiD675 containing *Cas5d, Cas6d, Cas7d*, and crRNA expression cassettes as shown in **Supplemental Fig. S6**. For PAM screening, the PAM screening reporter plasmid was constructed, assembling the sequence of the *lacI* gene, the *lacI* promoter to the 129th codon of the *lacZ* gene following the *ccdB* gene in pMW219 (Nippongene) to yield pPAM-ccdB. The PAM library fragment containing a target with a 4-nucleotide randomized PAM region was synthesized (IDT) and inserted in front of the *lacZ-ccdB* gene of pPAM-ccdB to yield pPAMlib-ccdB.

#### Mammalian vectors

Gene fragments corresponding to human codon-optimized *Cas* effector genes; *Cas3d, Cas5d, Cas6d, Cas7d*, and *Cas10d*, were synthesized with the SV40 nuclear localization signal (NLS) at their N-termini (gBlocks®) (IDT), assembled, and cloned into the pEFs vector (López-Perrote et al, 2016) separately to yield pEFs-HA-SV40NLS-Cas3d, pEFs-Strept-SV40NLS-Cas5d, pEFs-myc-SV40NLS-Cas6d, pEFs-FLAG-SV40NLS-Cas7d, pEFs-6xHis-SV40NLS-Cas10d. The tags fused to each Cas protein were as follows; HA-tag to Cas3d, Strep-tag to Cas5d, Myc -tag to Cas6d, FLAG-tag to Cas7d, and 6x His-tag to Cas10d. All *Cas* genes were combined into an all-in-one vector, pEFs-All, by fusing with sequences encoding 2A self-cleavage peptides. pEFs_Cas3d-Cas10d and pEFs_Cas6d-Cas5d-Cas7d expression vectors were also constructed by linking to the 2A self-cleavage peptide. To evaluate different promoters to express Cas5d and Cas6d, the human EFs promoter was replaced with the CAG promoter in pEFs-Strept-SV40NLS-Cas5d, and pEFs-Myc-SV40NLS-Cas6d to yield pCAG-Strept-SV40NLS-Cas5d or pCAG-Myc-SV40NLS-Cas6d. To construct the Cas-expressing vector with bipartite NLS (bpNLS), the DNA fragment (myc-bpNLS-Cas10d-bpNLS-6xHis) was first synthesized (IDT) and cloned into the pEFs vector, resulting in pEFs-Myc-bpNLS-Cas10d-bpNLS-6xHis vector. The Cas10d gene fragment in pEFs-Myc-bpNLS-Cas10d-bpNLS-6xHis vector was replaced by *Cas3d, Cas5d, Cas6d* and *Cas7d* gene fragments, respectively, to yield pEFs-Myc-bpNLS-Cas3d-bpNLS-6xHis, pEFs-Myc-bpNLS-Cas5d-bpNLS-6xHis, pEFs-Myc-bpNLS-Cas7d-bpNLS-6xHis, and pEFs-Myc-bpNLS-Cas10d-bpNLS-6xHis. To construct the mutated Cas10d (H177A) expression vector, the dCas10d (H177A) fragment was synthesized (IDT) and the wild-type Cas10d fragment pEFs-Myc-bpNLS-Cas10d-bpNLS-6xHis was replaced by dCas10d (H177A) to yield pEFs-myc-bpNLS-Cas (H177A)-bpNLS-6xHis. The Myc-bpNLS-Cas7d-bpNLS-6xHis fragment in pEFs-Myc-bpNLS-Cas7d-bpNLS was replaced by the 6xHis-myc-Cas7d-bpNLS fragment to yield pEFs-6xHis-myc-bpNLS-Cas7d-bpNLS. For crRNA expression, a DNA fragment containing a repeat-spacer-repeat sequence was cloned under the human U6 promoter in pEX-A2J1 (Eurofins Genomics) to yield pAEX-hU6crRNA. To construct pAEX-hU6crRNA_mature, two repeat sequences replaced the predicted processed repeat sequences. For insertion of gRNA sequence, two oligonucleotides containing a target sequence were annealed and cloned into the gRNA expression vector using Golden Gate cloning using the restriction enzyme BsaI (NEB).

#### Luc reporter assay plasmids

For the luc reporter assay, the NanoLUxxUC expression vector was constructed. First, NLUxxUC_Block1 and NLUxxUC_Block2 DNA fragments was synthesized (IDT). NLUxxUC_Block1 includes the first 351 bp of the NanoLUC gene sequence and Multi Cloning Site. An XbaI site was attached to the 5’ end of the NanoLUC gene. NLUxxUC_Block2 includes 465 bp of the 3’ end of the NanoLUC gene. An XhoI site was attached to the 3’ end of the NanoLUC gene. These fragments were assembled and cloned into the pCAG-EGxxFP vector obtained from addgene (#50716). Each split type NLUxxUC reporter was constructed by removing NLUxxUC_Block1 by XbaI and BamHI digestion, and the NLUxxUC_Block2 by XbaI and EcoRI digestion from pCAG-NLUxxUC vector, respectively. Each digested vector was assembled with a Multi Cloning Site, resulting in pCAG-NLUxxUC_Block1 and pCAG-NLUxxUC_Block2.

### Plasmid interference assay

*Escherichia coli* strain BL21-AI [F-*omp*T *hsd*SB (rB^-^mB^-^) *gal dcm ara*B∷ T7RNAP-*tet*A] (Thermo Fisher Scientific) was used in the plasmid interference assay. *E. coli* cells harboring pCmMaTiD675 were grown in LB medium supplemented with chloramphenicol (30 mg/ml). The PAM screening reporter plasmid library pPAMlib-ccdB was introduced into *E.coli* cells harboring pCmMaTiD675. After transformation, *E.coli* cells were pre-cultured in SOB liquid medium supplemented with 0.2% arabinose at 37°C for 2hr and plated onto LB agar medium supplemented with 0.2% arabinose, 0.4 mM IPTG, chloramphenicol (30 mg/mL) and kanamycin (25 mg/mL) at 37°C overnight. Plasmid DNA was extracted from surviving *E. coli* clones, and the PAM sequence was amplified with adapters for Illumina sequencing from extracted plasmid DNAs as templates. The 4-nt PAM regions from 300–400 reads of Illumina sequencing were analyzed. The experiments were repeated three times independently with similar results.

### Cell culture and transfection

Human embryonic kidney cell line 293T (HEK293T, RIKEN BRC) was cultured in Dulbecco’s modified Eagle’s Medium (DMEM) supplemented with 10% fetal bovine serum (Thermo Fisher Scientific), GlutalMAX™ Supplement (Thermo Fisher Scientific), 100 units/mL penicillin, and 100 μg/mL streptomycin in a 60 mm dish at 37°C with 5% CO_2_ incubation. HEK293T cells were seeded into 6-well plates (Corning) the day before transfection. Cells were transfected using TurboFect Transfection Reagent (Thermo Fisher Scientific) following the manufacturer’s recommended protocol. For each well of a 6-well plate, a total of 4 μg plasmids, extracted using NucleoSpin^**®**^ Plasmid Transfection-grade kit (Macherey-Nage), was used. Transfected cells were harvested after 48 hrs for mutation analysis.

### *In vitro* nuclease assay

*In vitro* nuclease assay was performed according to the protocol described in Sinkunas *et al*., (2011) with some modifications. M13mp18 single-strand DNA (NEB) was used as a substrate. Cas3d and Cas10d proteins were prepared by Unitech, Co. Briefly, Cas3d and Cas10d proteins were purified from HEK293T cells expressing each His-tagged Cas protein using Ni-NTA agarose (Qiagen) and a gel filtration column (Superdex 200 increase 10/300 GL columns). The Cas proteins were eluted in buffer (20 mM HEPES pH 7.5, 150 mM KCl, 1 mM DTT, 10% glycerol). The nuclease reaction was performed in buffer [10 mM HEPES pH 7.5, 75 mM KCl, 0.5 mM DTT, 5% glycerol, 2 mM ATP, 100 μM NiCl_2_, 100 μM CoCl_2_, 1 x Cut Smart Buffer (NEB), 4 nM M13mp18 ssDNA, 0.75 μM Cas protein] at 37 °C for 30 min, 1 hr and 2 hr, respectively. Reactions were stopped by adding chloroform followed by chloroform extraction. The aqueous phase was separated and mixed with Gel Loading Dye, Purple (6X) (NEB), then subjected to electrophoresis on a 1% agarose gel. The DNAs were visualized by staining with GelRed™ Nucleic Acid Gel Stain (Biotium). The experiments were repeated three times independently with similar results.

### Luc reporter assay

HEK293T cells were seeded into 96-well plates (Corning) the day before transfection at a density of 2.0 × 10^4^ cells/well. Cells were transfected using TurboFect Transfection Reagent (Thermo Fisher Scientific) following the manufacturer’s recommended protocol. For each well of a 96-well plate, a total of 200 ng plasmid DNAs including (1) pGL4.53 vector encoding *Fluc* gene (Promega), (2) pCAG-nLUxxUC vector interrupted with target DNA fragment, and (3) plasmid DNAs encoding TiD components were used. The NanoLuc and Fluc luciferase activities were measured 3 days after transfection using the Nano-Glo® Dual-Luciferase® Reporter Assay System (Promega). The firefly (Fluc) activity was used as an internal control. The NanoLuc/Fluc ratio was calculated for each sample and compared with the NanoLuc/Fluc ratio of the control sample, which was transfected with non-targeting gRNAs. The relative NanoLuc/Fluc activity was used to evaluate the gRNA activity. The experiments were repeated three times independently with similar results.

### Western blotting

HEK293T cells were transfected with 4 μg of plasmid DNA in 6-well plates. At 2d post-transfection, total proteins were extracted from HEK293T cells using RIPA Lysis and Extraction Buffer (Thermo Scientific) supplemented with Protease Inhibitor Cocktail for Use with Mammalian Cell and Tissue Extracts (Nakalai Tesque) according to the manufacturer’s protocol. For the isolation of nuclear and cytoplasmic proteins, NE-PER™ Nuclear and Cytoplasmic Extraction Reagents (Thermo Fisher Scientific) was used.

The extracted proteins were quantified using a Pierce™ BCA Protein Assay Kit (Thermo Fisher Scientific). The samples were mixed with 4 x Laemmli Sample buffer (Bio-Rad) and 2.5% β-mercaptoethanol, followed by heat treatment at 95°C for 5 min. The denatured proteins were loaded into 12% SDS-PAGE gel with Tris-Glycine-SDS buffer [0.25M Tris, 1.92M Glycine, 1% (w/v) SDS] and separated by SDS-PAGE for 60 min at 150V. The proteins were transferred to Immobilon-P Polyvinylidene fluoride (PVDF) membranes (Millipore) in Tris-Glycine-SDS buffer containing 20% methanol for 2h at 50V. The blot was washed by TTBS (25mM Tris, 137 nM NaCl, 2.68 nM KCl) for 5 min 3 times and blocked at room temperature in Blocking One (Nacalai Tesque) for 60 min. The primary antibody reactions were performed at room temperature for 60 min. After the membranes were washed by TTBS for 5 min 3 times, the secondary antibody, Anti-Mouse IgG (H+L), HRP Conjugate (Promega, 1:10000 in Blocking one), was added to the membrane, After incubation for 60 min at room temperature, the membrane was washed with TTBS 5 min 3 times. Signal was detected using SuperSignal™ West Pico PLUS Chemiluminescent Substrate (Thermo Fisher Scientific) according to the protocol provided by the manufacturer. Images were captured using an ImageQuant LAS4000 mini (GE Healthcare Bioscience). Primary antibodies used in this study are as follows: Anti-DDDDK-tag mAb (1:10,000) (MBL, Japan), Anti-HA-tag mAb (1:10,000) (MBL), Anti-His-tag mAb (1:10,000) (MBL), Anti-Myc-tag mAb (1:10,000) (MBL), Anti-Strep-tag mAb (1:1000) (MBL), Anti-*β*-actin mAb (1:10000) (MBL). The experiments were repeated times independently with similar results.

### Immunoprecipitation

HEK293T cells were transfected with 1 μg of each Cas expression vector (pEFs-FLAG-SV40NLS-Cas7d, pEFs-Myc-bpNLS-Cas3d-bpNLS-6xHis, pEFs-Myc-bpNLS-Cas5d-bpNLS-6xHis, pEFs-Myc-bpNLS-Cas6d-bpNLS-6xHis, pEFs-Myc-bpNLS-Cas7d-bpNLS-6xHis, pEFs-Myc-bpNLS-Cas10d-bpNLS-6xHis) and gRNA expression vector using TurboFect Transfection Reagent (Thermo Fisher Scientific) on a 60 mm dish. Protein extraction was performed according to the protocol described in Katoh et al., (2015) at 48h post-transfection. Briefly, the medium was replaced with Lysis buffer (20 mM HEPES, 150 mM NaCl, 0.1% (w/v) Triton X-100, 10% (w/v) Glycerol) containing Protease Inhibitor Cocktail for Use with Mammalian Cell and Tissue Extracts (Nacalai Tesque) and incubated for 5 min on ice. The cell lysates were mixed by pipetting and transferred to 1.5 ml tubes, then incubated for 15 min on ice. Purification of Cascade complex containing FLAG-NLS-Cas7d protein was performed using DDDDK-tagged Protein Magnetic Purification Kit (MBL) according to the protocol provided by the manufacturer. The resulting elutes were analyzed by SDS-PAGE and western blotting using following antibodies: Anti-His-tag mAb (1:10,000) (MBL), Anti-DDDDK-tag mAb (1:10,000) (MBL), Anti-*β*-actin mAb (1:10,000) (MBL), Anti-Mouse IgG (H+L), HRP Conjugate (1:10,000) (Promega). The experiments were repeated three times independently with similar results.

### DNA deletion analysis by long-range PCR

To detect DNA deletion in HEK293T cells, long-range PCR and cloning of a pool of long PCR products was performed. At first, genomic DNA was extracted from HEK293T cells using Geno Plus (TM) Genomic DNA Extraction Miniprep System (Viogene-BioTek). Then, nested PCR was performed to amplify the long-range DNA region specifically. At first, the extracted genome DNA was used as a template for the amplification of target DNA region using several kinds of specific-primer sets for longrange PCR (**Supplemental TableS4**), which were designed to amplify target DNA region of various lengths (3.5 kb to 25 kb). The first PCR reactions were performed using KOD ONE Master Mix (TOYOBO) under the following conditions: 35 cycles of 10 s at 98°C, 5 s at 60°C and 50 s (amplicon: 15–20 kb) or 150 s (amplicon: 10–15 kb) or 200 s (amplicon: <10 kb) at 68°C). Then, the PCR products were diluted 100–10,000 times and used as a template for the nested PCR. The nested PCR was also performed under same conditions mentioned above. The PCR products were separated by electrophoresis on 1% agarose gel and visualized by staining with GelRed™ Nucleic Acid Gel Stain (Biotium). Nested PCR products were pooled and purified using Monofas^®^ DNA purification Kit I (GL Sciences). The purified mixture of PCR products was cloned into pMD20-T vector using Mighty TA-cloning Kit (TaKaRa). The 119 clones for *AAVS*-*tid* GTC_70-107 (+) and 20 clones for *hEMX1-tid* GTT_9 (–) were picked up and Sanger sequenced using M13 Uni and M13 RV primers. Sanger sequencing results were analyzed using BLATN search and ClustalW program to identify the DNA deletions.

### Mutation analyses in short-range PCR products

To evaluate mutations introduced in transfected human cells, a region of about 400 bp surrounding the target locus of gRNA was amplified by short-range PCR using a PCR kit as described above. In HMA, the PCR fragments were analyzed directly using a microchip electrophoresis system with MCE202 MultiNA (Shimazu). PCR amplicons were also cloned into the TA cloning vector (TaKaRa) to determine their sequences by the Sanger method. All primers used for short-range PCRs used in the mutation analyses are listed in **Supplemental Table S3**.

## SUPPLEMENTAL INFORMATION

Supplemental Information can be found online at https://doi.org/xxxxxxxxxx

## ACKNOWLEDGEMENTS

We thank M. Fukuhara, and E. Niimi for their technical assistance. This research was supported by the New Energy and Industrial Technology Development Organization (NEDO; to K.O.). This work was also supported by Program on Open Innovation Platform with Enterprises, Research Institute and Academia (OPERA, funding agency: Japan Science and Technology Agency; to K.O.).

## AUTHOR CONTRIBUTION STATEMENTS

K.O. and Y.O designed and wrote the manuscript with help from all authors. N. W. designed and performed experiments using animal cells. E. M. performed experiments using bacterial and animal cells.

## DECLARATION OF INTERESTS

Two patent applications have been filed relating to the data presented. The authors have no conflict interest to declare.

## Notes

#### Summary of Updates

The title has been changed.

