## Supplemental materials for "Genome editing in mammals using CRISPR type I-D nuclease"

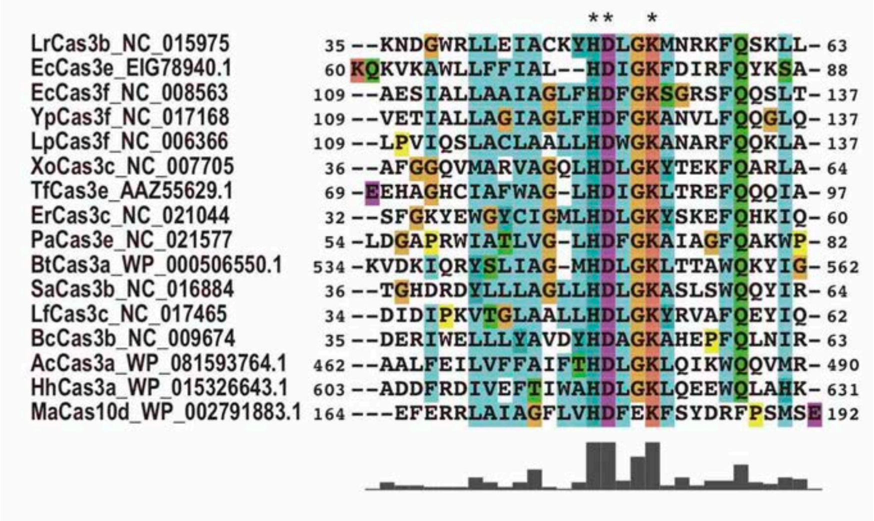


**Supplemental Fig. S1. Amino acid sequence alignment of the HD domain in Cas3 and Cas10d.**

Multiple alignment was generated by ClustalX using the Cas3 HD domains from various bacteria and a Cas10d HD domain from *Microcystis aeruginosa*. MaCas10d; *M. aeruginosa* (WP_002791883.1), AcCas3a; *Anabaena cylindrica* (WP_081593764.1), BtCas3a; *Bacillus thuringiensis* (WP_000506550.1), HhCas3a *Halobacteroides halobius* (WP_015326643.1), BcCas3b; *Bacillus cytotoxicus* (NC_009674), LrCas3b; *Lactobacillus ruminis* (NC_015975), SaCas3b; *Sulfobacillus acidophilus* (NC_016884), ErCas3c; *Eubacterium rectale* (NC_021044), LfCas3c; *Lactobacillus fermentum* (NC_017465), XoCas3c; *Xanthomonas oryzae* (NC_007705), EcCas3e; *Escherichia coli* (EIG78940.1), PaCas3e; *Pseudomonas aeruginosa* (NC_021577), TfCas3e; *Thermobifida fusca* (AAZ55629.1), EcCas3f; *Escherichia coli* (NC_008563), LpCas3f; *Legionella pneumophila* (NC_006366), YpCas3f; *Yersinia pestis* (NC_017168).

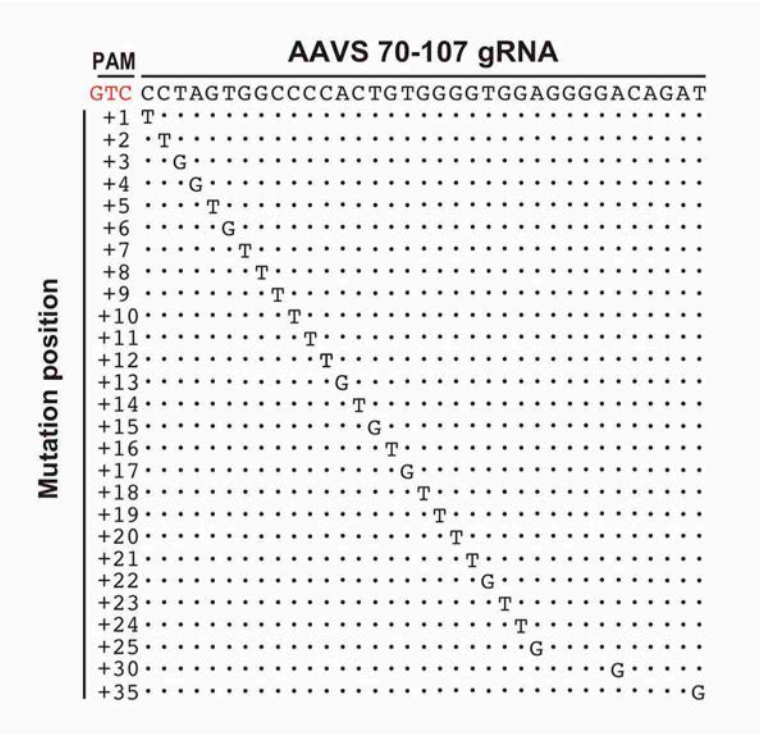


**Supplemental Fig. S2. Evaluation of off-target effects in the TiD gRNA.**
The critical nucleotides in the target gRNA sequence for genome editing activity of TiD were evaluated using the luc reporter assay in human HEK293T cells using the split-type vectors (**Fig. 2, Supplemental Method**). Variation of the gRNA sequences of *AAVS* GTC_70-107(+) were indicated.

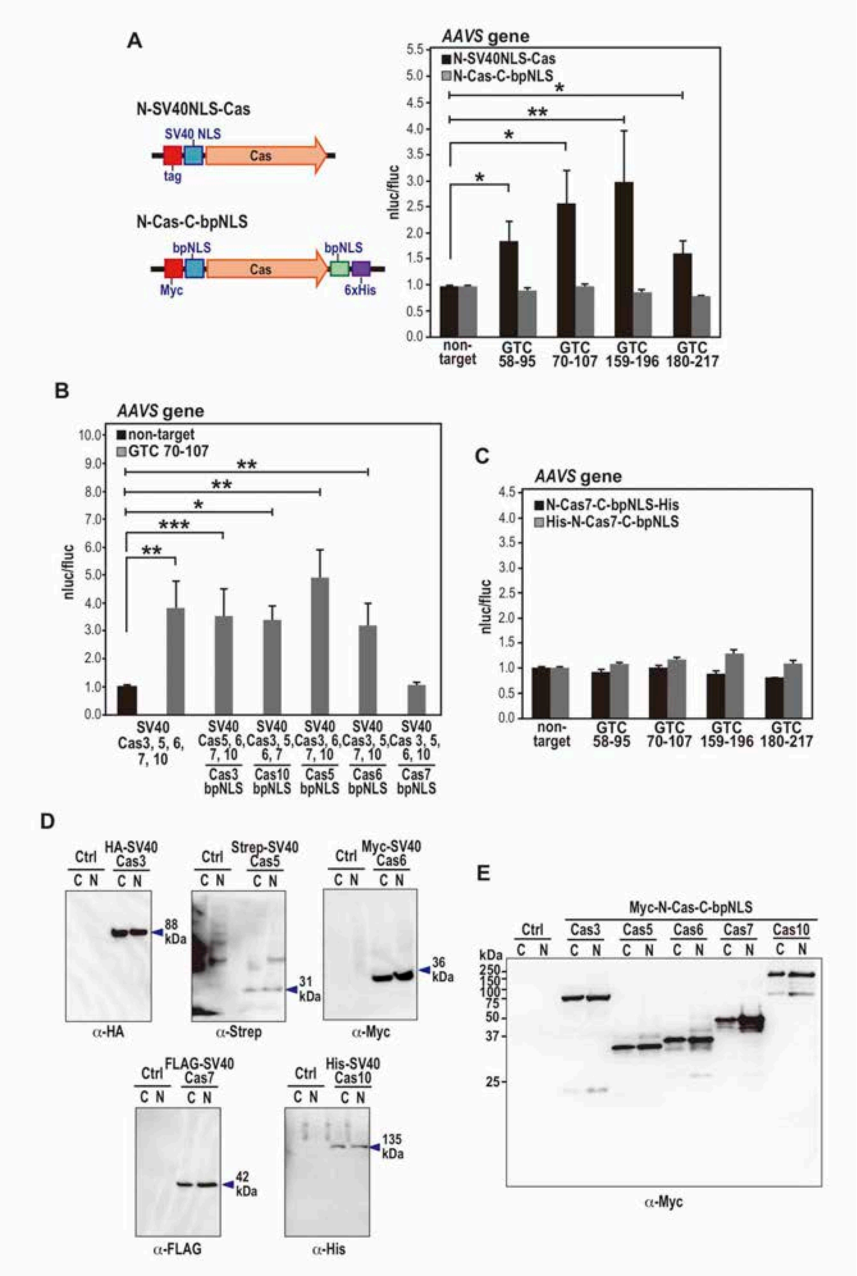

**Fig. S3. Effect of nuclear localization signals of Cas proteins in TiD activity.**

**(A)** Left; schematic structure of the Cas expression vector cassettes. N-SV40NLS-Cas; SV40NLS on N-ter with different tags in Cas3d, Cas5d, Cas6d, Cas7d, and Cas10d, respectively. N- Cas-C-bpNLS; bpNLSs on both N-ter with Myc-tag and C-ter with 6xHis-tag in Cas3d, Cas5d, Cas6d, Cas7d, and Cas10d, respectively. Right; Effect of the NLS of Cas proteins in the TiD activity to the various gRNAs targets for *AAVS* determined by the luc reporter assay in human HEK293T cells. Data are means ± S.E. of independent experiments (N=4). *P<0.1 and **P<0.05 are determined by Student's t tests.

**(B)** Left; bpNLS functioned effectively on both N-ter and C-ter in Cas3d, Cas5d, Cas6d, Cas10d as SV40NLS on N-ter in each Cas, however, bpNLS attached in Cas7d disrupted TiD activity.

The data suggests that the Cas7 inactivation affected the repression of TiD activity in **a** (right). Data are means ± S.E. of independent experiments (N=4). *$P<0.01$, **$P<0.05$, and ***$P<0.07$ are determined by Student's t tests.

**(C, D)** Protein blot analysis of the Cas-NLS proteins extracted from human HEK293T cells used in the luc reporter assay. C; cytosol fraction, N; nuclear fraction.

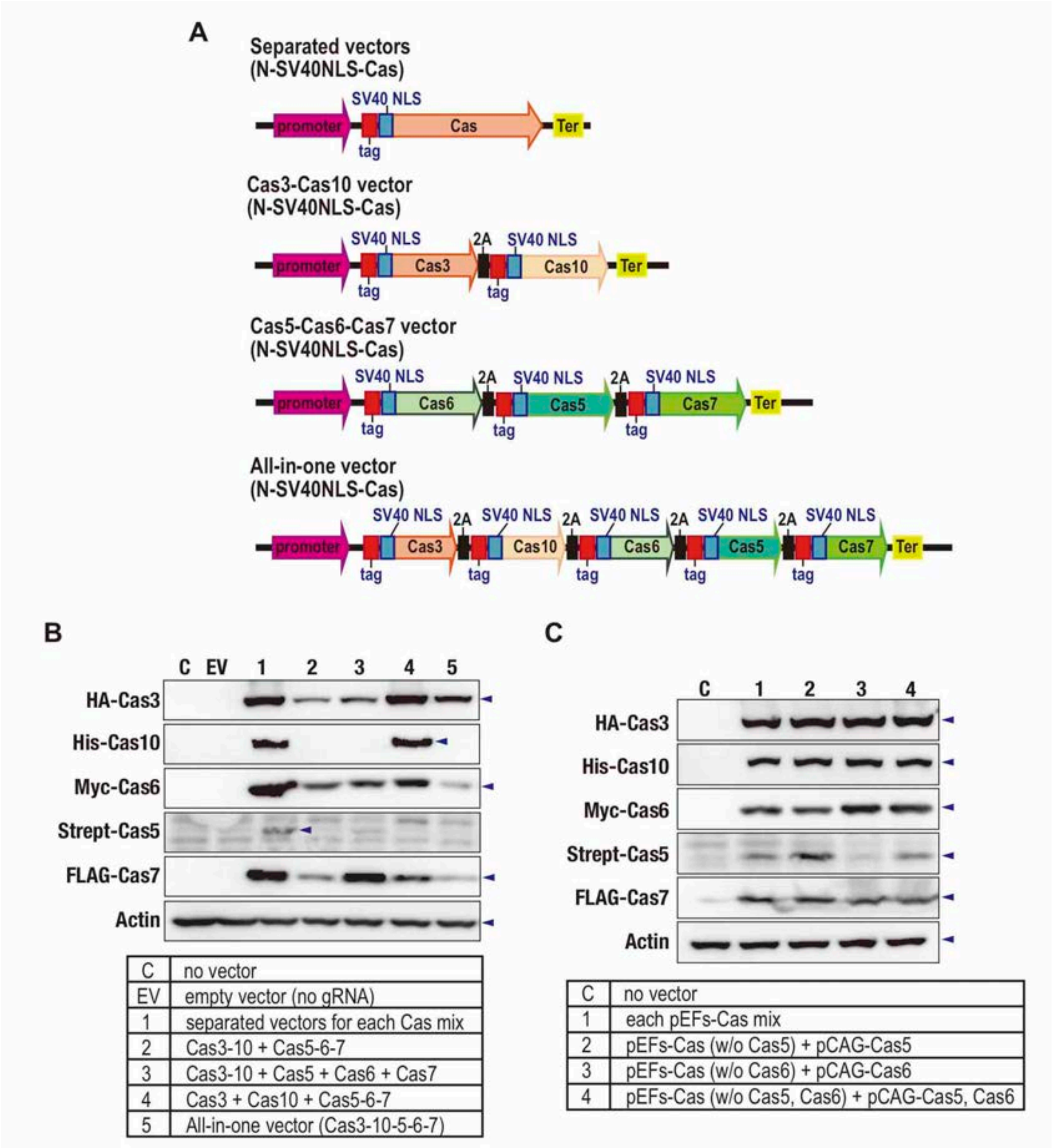


**Fig. S4. Optimization of the TiD expression vector in human HEK293T cells.**
**(A)** Schematic structures of the Cas expression vector cassettes using genome editing of human HEK293T cells. Separated each Cas vectors, and 2 sets type (Cas3-Cas10 and Cas5-Cas6-Cas7), and all-in-one type of vectors were used to compare the expression levels in **b, c**. Cas3 and Cas10, or Cas5, Cas6, and Cas7 in two vectors and all Cas proteins in all-in-one vector were fused via 2A self-cleaving peptide to generate the single transcriptional products and express simultaneously. N-SV40NLS-Cas; SV40NLS on N-ter with different tags in Cas3d, Cas5d, Cas6d, Cas7d, and Cas10d, respectively, 2A; self-cleaving peptide.
**(B)** Expression levels of Cas proteins in the HEK293T cells. 1; The Cas expression cassettes were separated in the individual vectors. 2-4; Cas3 and Cas10, or Cas5, Cas6, and Cas7 were inserted in the same vectors and transfected to HEK293T cells as the separated-type vectors. 5;

all-in-one vector for all Cas expression cassettes. The expression levels were detected by the western blot using the specific antibody for each tag fused to the Cas.

**(C)** Optimization of the Cas5 and Cas6 expression levels using the different promoter in the HEK293T cells. The expression levels were detected by the western blot using the specific antibody for each tag fused to the Cas. The *elongation Factor 1*α promotor in the pEF vector and the *chicken B actin* promoter in the pCAG were used for the Cas expression, respectively.

**A**

*AAVS-tid* GTC_70-107(+)

| large deletion (bp) | clone numbers | (%) |
|---|---|---|
| Δ18443 (-7784, +10659, 2272 bp insertion) | 1 | (1.5) |
| Δ18443 (-7775, +10673, MH* 5 bp) | 2 | (2.9) |
| Δ17902 (-7366, +10536, 87 bp insertion) | 18 | (26.5) |
| Δ17833 (-7297, +10536, 20 bp insertion) | 1 | (1.5) |
| Δ17817 (-7366, +10451, MH 31bp) | 2 | (2.9) |
| Δ17815 (-7298, +10517, MH 3bp) | 1 | (1.5) |
| Δ17218 (-6628, +10590, MH 1bp) | 2 | (2.9) |
| Δ7008 (-3610, +3401, MH 3 bp) | 1 | (1.5) |
| Δ5494 (-3039, +2455) | 12 | (17.6) |
| Δ5235 (-2671, +2565, MH 1bp) | 11 | (16.2) |
| Δ5057 (-3644, +1416, MH 3bp) | 2 | (2.9) |
| Δ4531 (-2309, +2222, 1bp insertion) | 2 | (2.9) |
| Δ3994 (-3607, +389, MH 2 bp) | 3 | (4.4) |
| Δ3820 (-321, +3504, MH 5 bp) | 3 | (4.4) |
| Δ3185 (-461, +2724) | 1 | (1.5) |
| Δ2589 (-852, +1738, MH 1 bp) | 6 | (8.8) |

*MH; micro homology

**B**

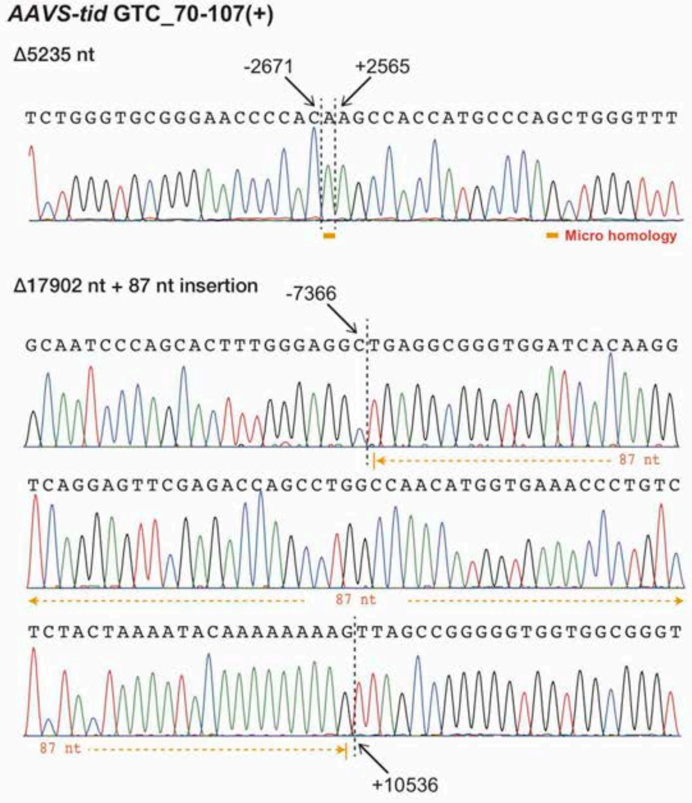


**Supplemental Fig. S5. The detection of long-range deletion mutations in the *hEMX1* gene induced by the CRISPR TiD.**

**(A)**. The *hEMX1* gene structure, gRNA positions, and the various primer sets (arrows) to amplify the mutation.

**(B)** The PCR amplified fragments separated on agarose gels. Numbers show the primer sets shown in the gene structures.

**(C)** The large deletion mutations analyzed by the Sanger sequencing of the cloned DNA from the CRISPR TiD infected cells. The nucleotide positions from the PAM were indicated on the sequence.

**Supplemental Fig. S6 Plasmid DNAs used in this study.**

**List of Plasmid DNAs**

(1) pCmMa567Δ10
(2) pCmMa567
(3) pPAMlib-ccdB
(4) pEFs-HA-SV40NLS-Cas3d
(5) pEFs-Strept-SV40NLS-Cas5d
(6) pEFs-myc-SV40NLS-Cas6d
(7) pEFs-FLAG-SV40NLS-Cas7d
(8) pEFs-6xHis-SV40NLS-Cas10d
(9) pEFs-All
(10) pEFs-Cas3d-Cas10d
(11) pEFs-Cas5d-Cas6d-Cas7d
(12) pCAG-Strept-SV40NLS-Cas5d
(13) pCAG-Myc-SV40NLS-Cas6d
(14) pEFs-Myc-bpNLS-Cas3d-bpNLS-6xHis
(15) pEFs-Myc-bpNLS-Cas5d-bpNLS-6xHis
(16) pEFs-Myc-bpNLS-Cas6d-bpNLS-6xHis
(17) pEFs-Myc-bpNLS-Cas7d-bpNLS-6xHis
(18) pEFs-Myc-bpNLS-Cas10d-bpNLS-6xHis
(19) pEFs-Myc-bpNLS-Cas10d(H177A)-bpNLS-6xHis
(20) pEFs-6xHis-Myc-Cas7d-bpNLS
(21) pAEX-hU6crRNA
(22) pAEX-hU6crRNA_mature
(23) pCAG-nLUxxUC
(24) pCAG-nLUxxUC_Block1_MCS
(25) pCAG-nLUxxUC_MCS_Block2

**Plasmid name: pCmMa567∆10**

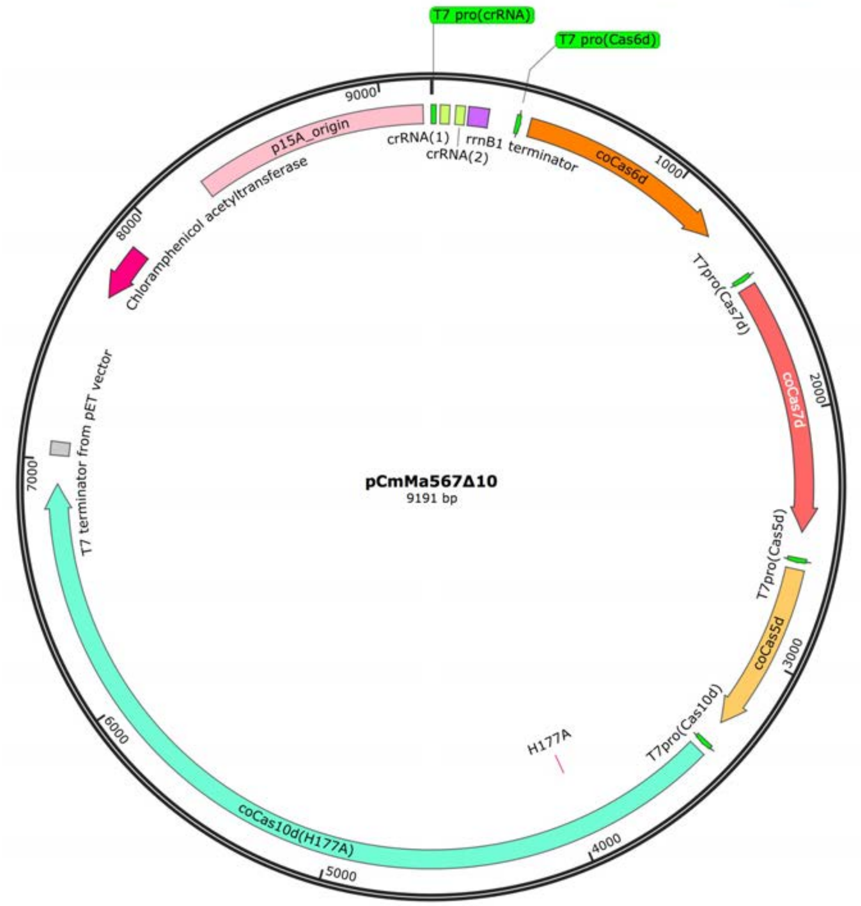


1..19: T7 promoter
36..72: crRNA(1)
96..132: crRNA(2)
146..225: rrnB1 terminator
335..353: T7 promoter
390..1223: coCas6d
1430..1448T7 promoter
1475..2476: coCas7d
2579..2597: T7 promoter
2624..3298: coCas5d
3390..3408: T7 promoter
3435..6905: coCas10d
3963..3965: H177A
7017..7069: T7 terminator from pET vector
7671..7886 (complement): Chloramphenicol acetyltransferase gene
8248..9160: p15A origin

**Plasmid name: pCmMa567**

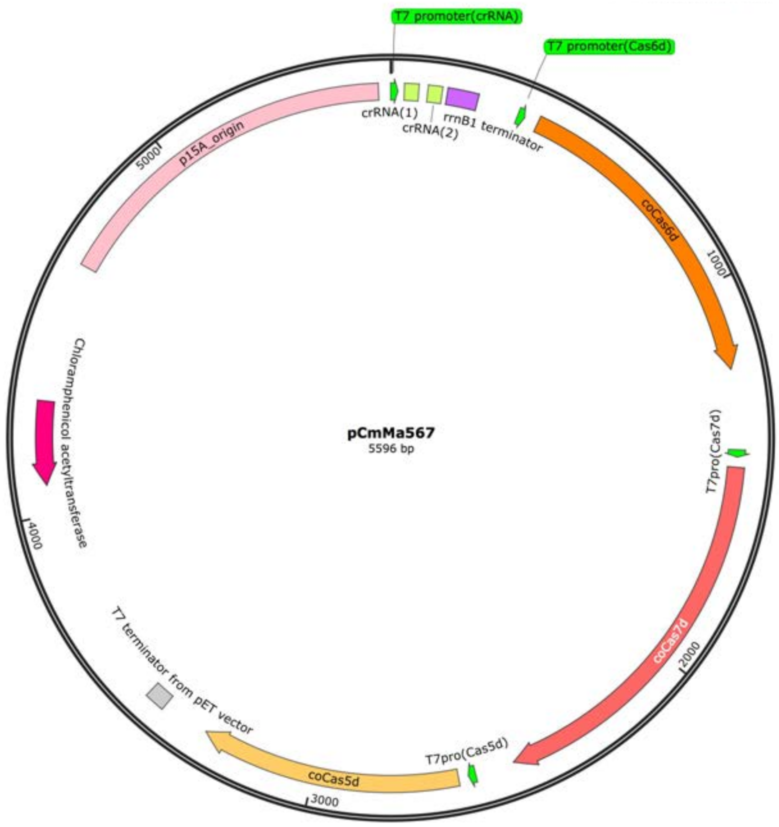


1..19: T7 promoter
36..72: crRNA(1)
96..132: crRNA(2)
146..225: rrnB1 terminator
335..353: T7 promoter
390..1223: coCas6d
1430..1448T7 promoter
1475..2476: coCas7d
2579..2597: T7 promoter
2624..3298: coCas5d
3422..3474: T7 terminator from pET vector
4076..4291 (complement): Chloramphenicol acetyltransferase gene
4653..5565: p15A origin

**Plasmid name: pPAMlib-ccdB**

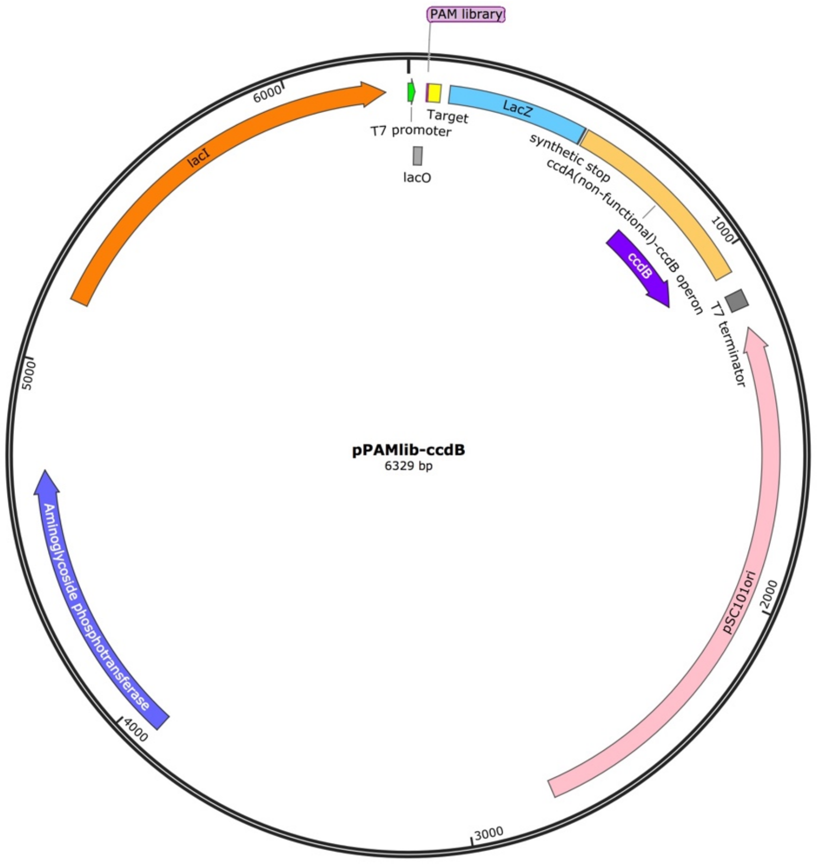


1..19: T7 promoter
19..46: lacO
56..90: Traget, 35nt
52..55: PAM library, 4nt
116..504: LacZ gene, #1 to #386 with synthetic stop codon
511..1063: ccdA(non-functional)-ccdB operon
758..1063: ccdB
1118..1164: T7 terminator
1221..2756(complement): pSC101ori
3912..4706: Aminoglycoside phosphotransferase
5185..6267: lacI

**Plasmid name: pEFs-HA-SV40NLS-Cas3d**

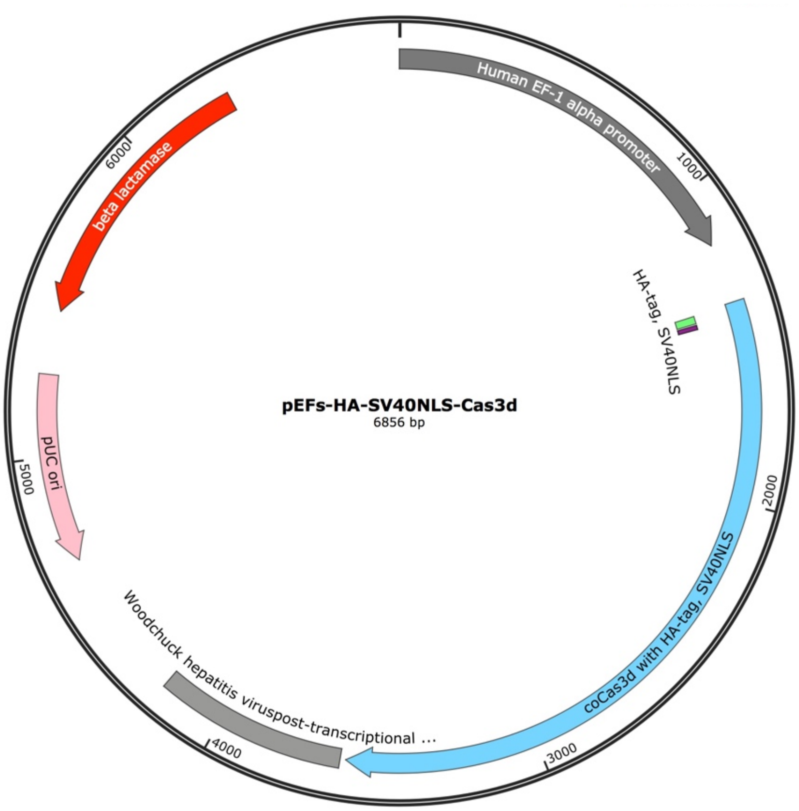


1..1182: Human EF-1 alpha promoter
1370..3595: coCas3d with HA-tag, SV40NLS
1376..1402: HA-tag
1409..1423: SV40NLS
3613..4201: Woodchuck hepatitis virus post-transcriptionalregulatory element (WPRE)
5456..6316(complement): beta lactamase
4668..5256(complement): pUC ori

**Plasmid name: pEFs-Strept-SV40NLS-Cas5d**

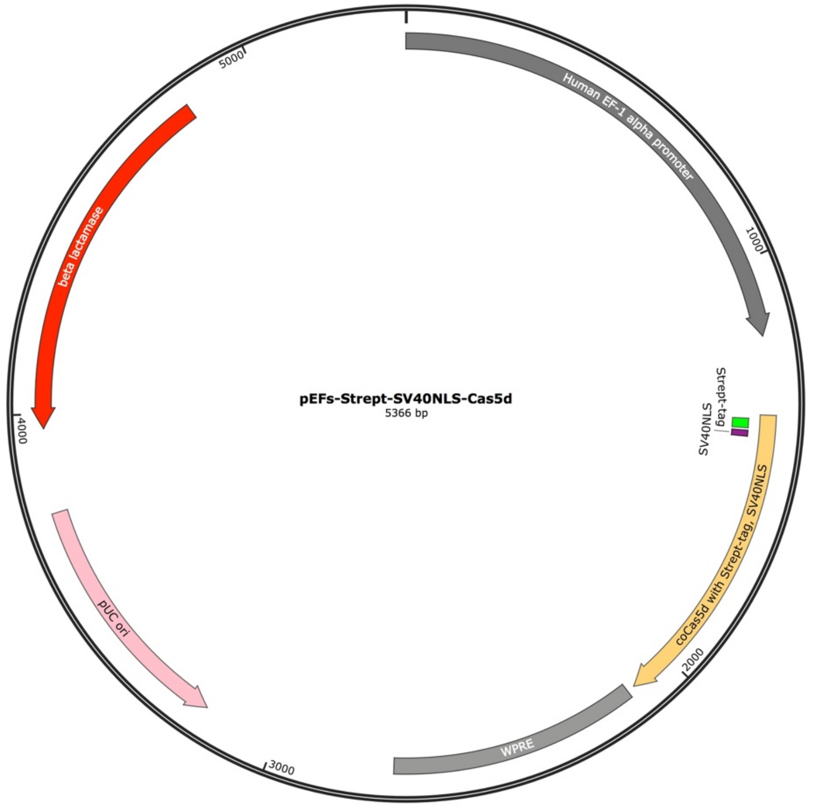


1..1182: Human EF-1 alpha promoter
1370..2104: coCas5d with Strept-tag, SV40NLS
1379..1402: Strept-tag
1409..1423: SV40NLS
2123..2711: Woodchuck hepatitis virus posttranscriptional regulatory element (WPRE)
3178..3766(complement): pUC ori
3966..4826(complement): beta lactamase

**Plasmid name: pEFs-myc-SV40NLS-Cas6d**

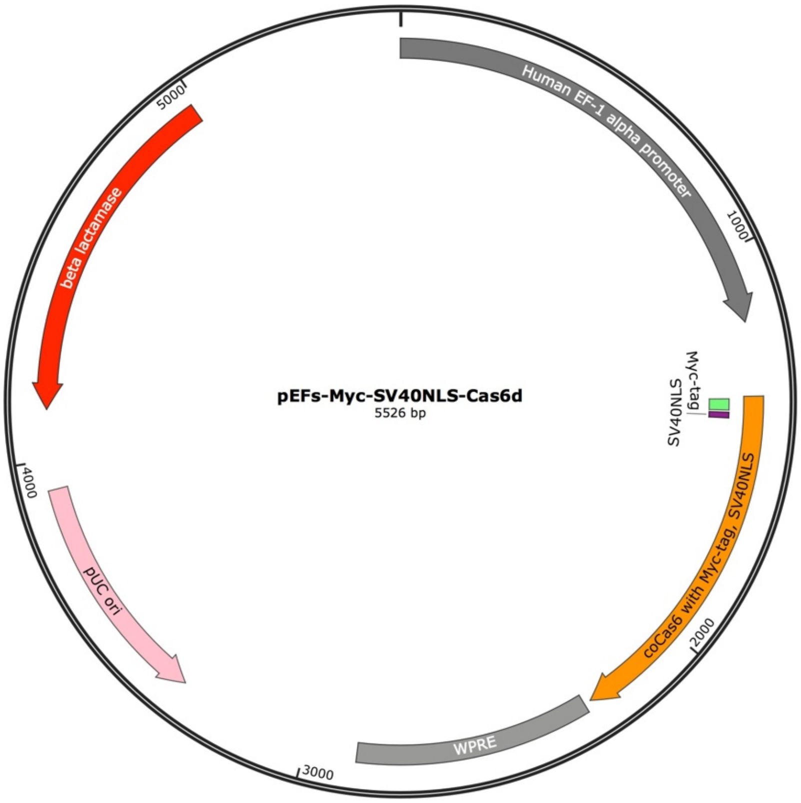


1..1182: Human EF-1 alpha promoter
1370..2104: coCas5d with Strept-tag, SV40NLS
1375..1404: Myc-tag
1411..1425: SV40NLS
2283..2871: Woodchuck hepatitis virus posttranscriptional regulatory element (WPRE)
3338..3926(complement): pUC ori
4126..4986(complement): beta lactamase

**Plasmid name: pEFs-FLAG-SV40NLS-Cas7d**

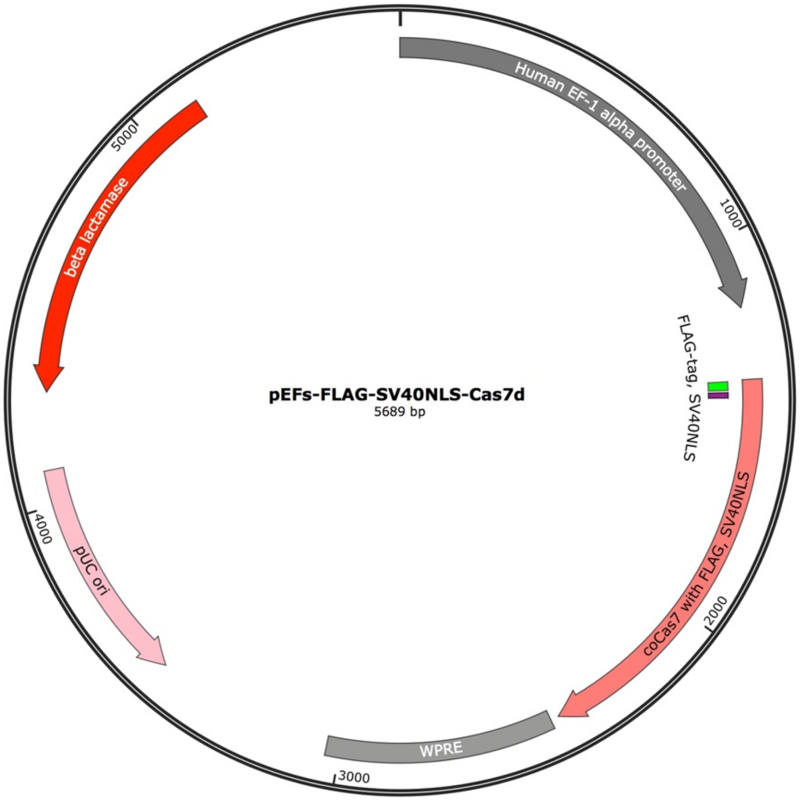


1..1182: Human EF-1 alpha promoter
1369..2424: coCas7d with FLAG-tag, SV40NLS
1372..1395: FLAG-tag
1402..1416: SV40NLS
2446..3034: Woodchuck hepatitis virus posttranscriptional regulatory element (WPRE)
3501..4089(complement): pUC ori
4289..5149(complement): beta lactamase

**Plasmid name: pEFs-6xHis-SV40NLS-Cas10d**

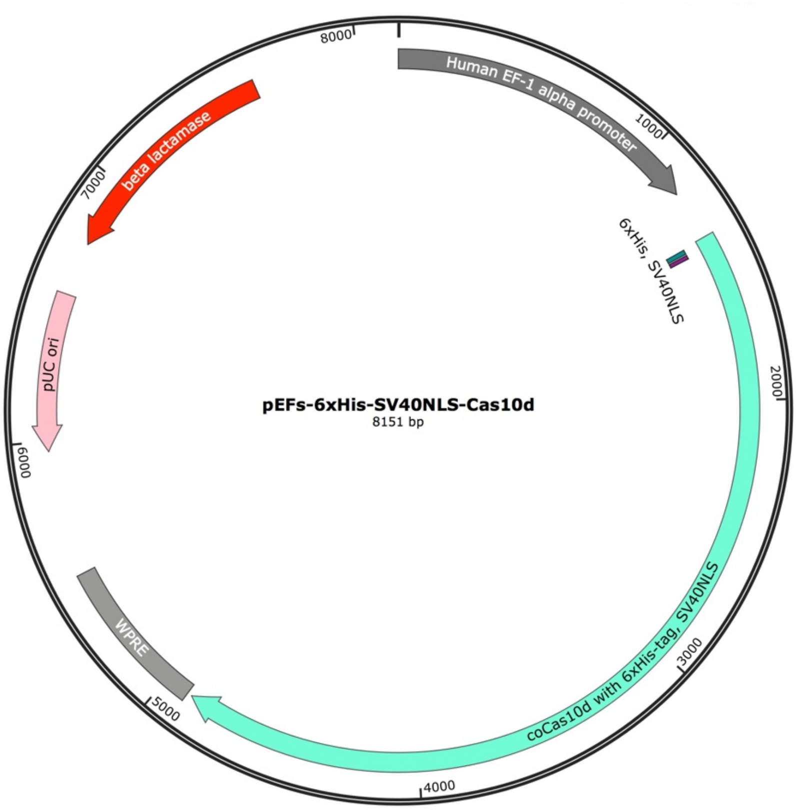


1..1182: Human EF-1 alpha promoter
1369..4890: coCas10d with 6xHis-tag, SV40NLS
1375..1392: 6xHis-tag
1399..1413: SV40NLS
4908..5496: Woodchuck hepatitis virus posttranscriptional regulatory element (WPRE)
5963..6551(complement): pUC ori
6751..7611(complement): beta lactamase

**Plasmid name: pEFs-All**

1..1182: Human EF-1 alpha promoter

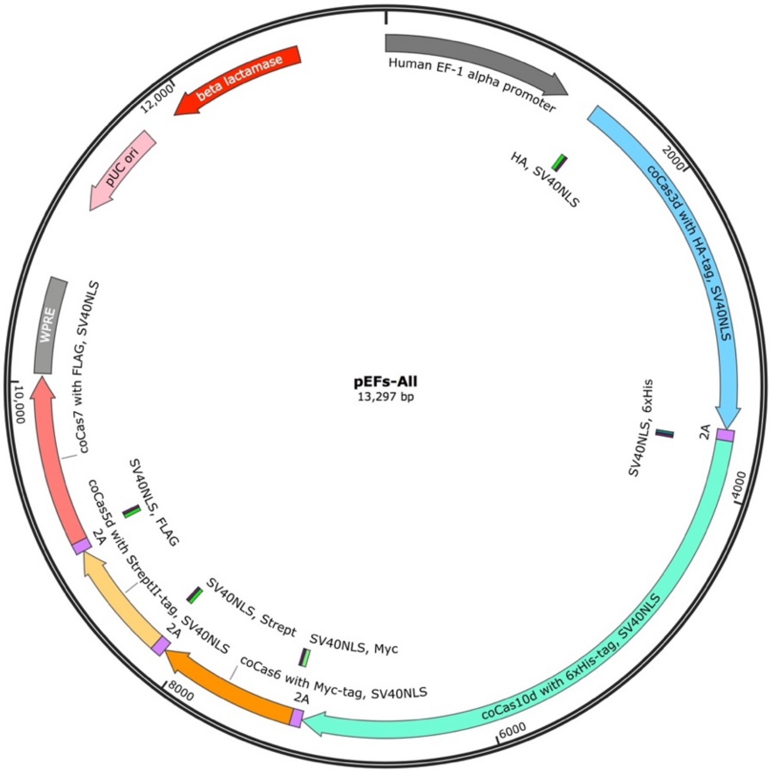


1370..3592: coCas3d with HA-tag, SV40NLS
1376..1402: HA-tag
1409..1423: SV40NLS
3593..3658: 2A self-cleavage peptide
3659..7171: coCas10d with 6xHis-tag, SV40NLS
3659..3676: 6xHis-tag
3683..3697: SV40NLS
7172..7237: 2A self-cleavage peptide
7238..8125: coCas6d with Myc-tag, SV40NLS
7238..7267: Myc-tag
7274..7288: SV40NLS
8126..8194: 2A self-cleavage peptide
8195..8917: coCas5d with Strept-tag, SV40NLS
8195..8218: Strept-tag
8225..8239: SV40NLS
8918..8983: 2A self-cleavage peptide
8984..10036: coCas7d with FLAG-tag, SV40NLS
8984..9007: FLAGs-tag
9014..9028: SV40NLS
10054..10642: Woodchuck hepatitis virus posttranscriptional regulatory element (WPRE)
11109..11697(complement): pUC ori
11897..12757(complement): beta lactamase

**Plasmid name: pEFs-Cas3d-Cas10d**

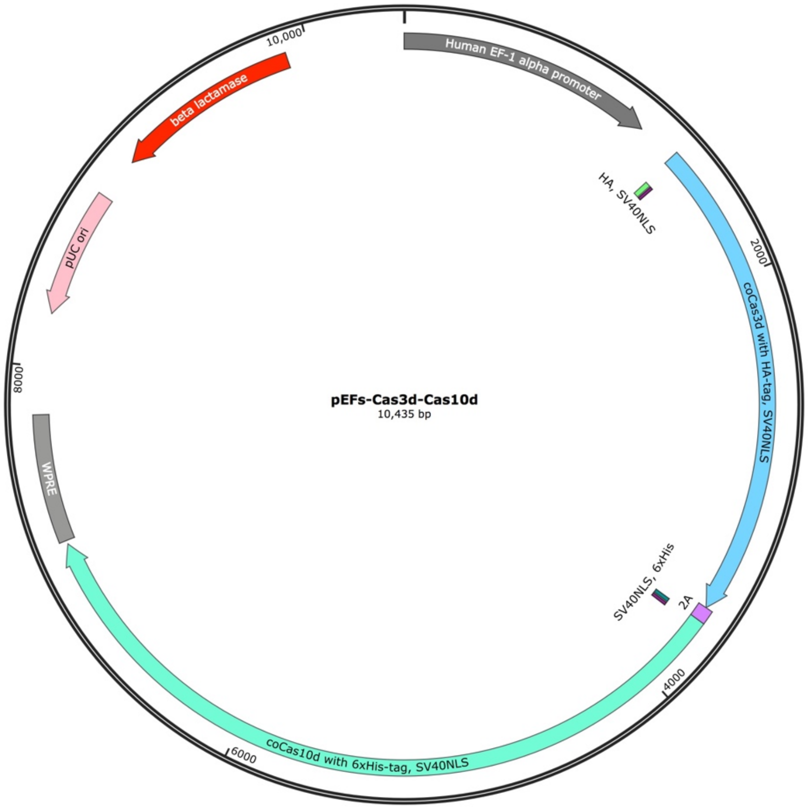


1..1182: Human EF-1 alpha promoter
1370..3592: coCas3d with HA-tag, SV40NLS
1376..1402: HA-tag
1409..1423: SV40NLS
3593..3658: 2A self-cleavage peptide
3659..7174: coCas10d with 6xHis-tag, SV40NLS
3659..3676: 6xHis-tag
3683..3697: SV40NLS
7192..7780: Woodchuck hepatitis virus posttranscriptional regulatory element (WPRE)
8247..8835(complement): pUC ori
9035..9895(complement): beta lactamase

**Plasmid name: pEFs-Cas5d-Cas6d-Cas7d**

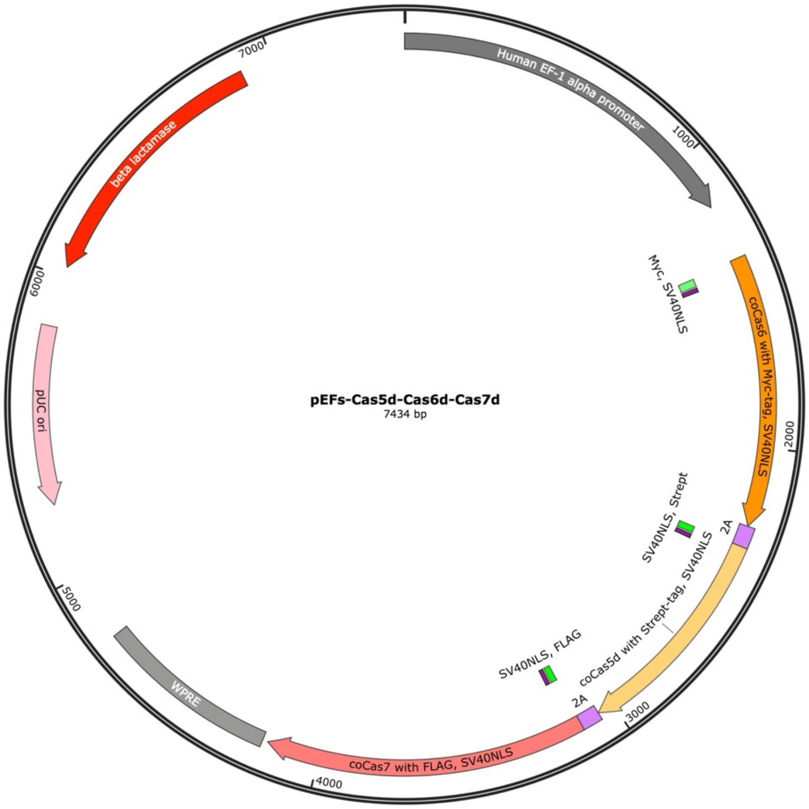


1..1182: Human EF-1 alpha promoter
1369..2262: coCas6d with Myc-tag, SV40NLS
1375..1404: Myc-tag
1411..1425: SV40NLS
2263..2331: 2A self-cleavage peptide
2332..3054: coCas5d with Strept-tag, SV40NLS
2332..2355: Strept-tag
2362..2376: SV40NLS
3055..3120: 2A self-cleavage peptide
3121..4173: coCas7d with FLAG-tag, SV40NLS
3121..3144: FLAGs-tag
3151..3165: SV40NLS
4191..4779: Woodchuck hepatitis virus posttranscriptional regulatory element (WPRE)
5246..5834(complement): pUC ori
6034..6894(complement): beta lactamase

**Plasmid name: pCAG_Strept-SV40NLS-Cas5d**

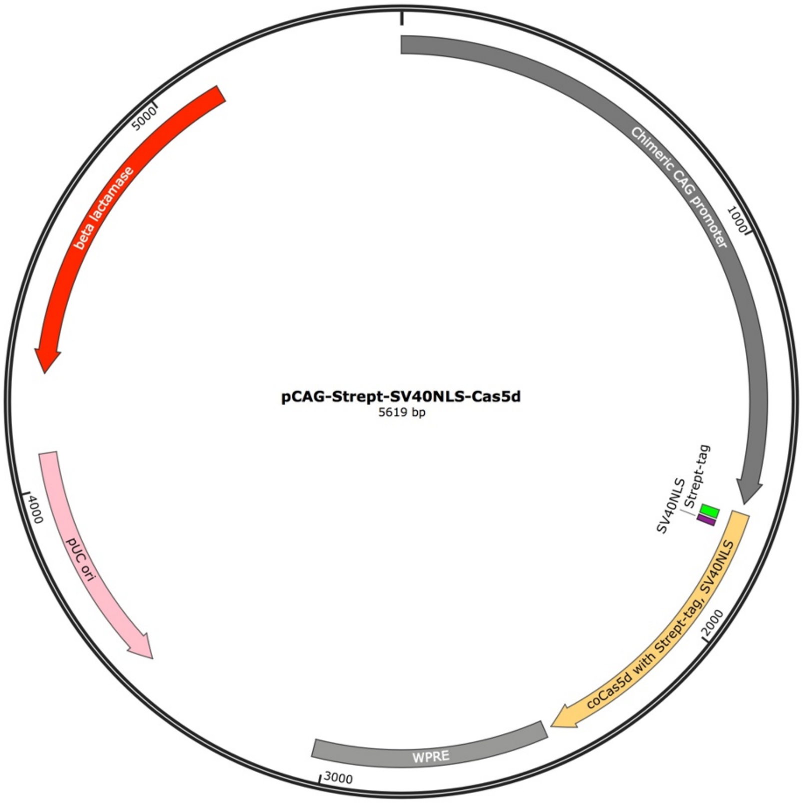


1..1665: Chimeric CAG promoter
1690..2424: coCas5d with Strept-tag, SV40NLS
1699..1722: Strept-tag
1729..1743: SV40NLS
2443..3031: Woodchuck hepatitis virus posttranscriptional regulatory element (WPRE)
3498..4086(complement): pUC ori
4286..5146(complement): beta lactamase

**Plasmid name: pCAG-Myc-SV40NLS-Cas6d**

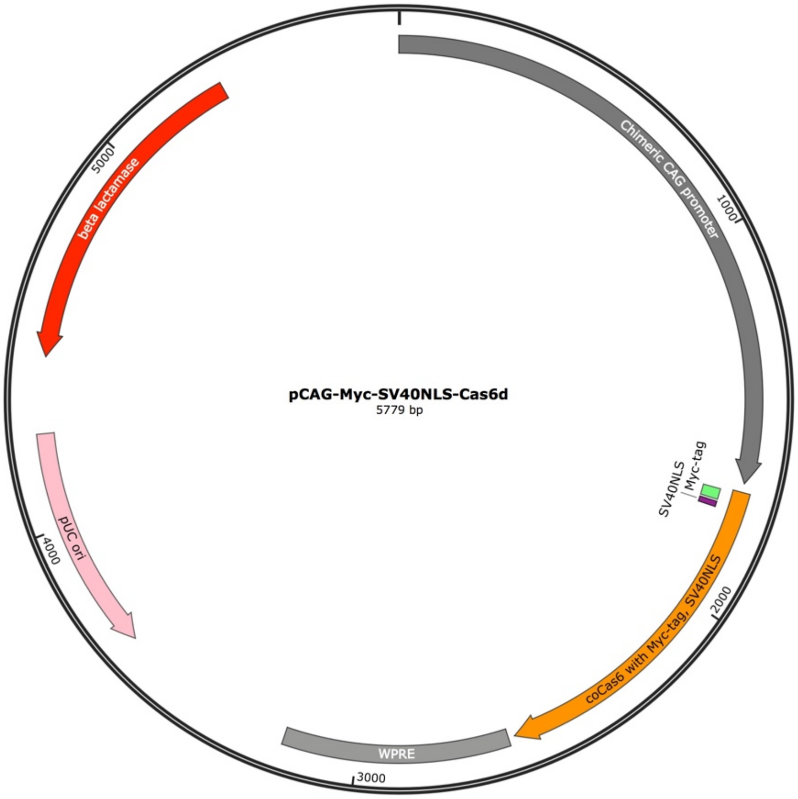


1..1665: Chimeric CAG promoter
1689..2585: coCas6d with Myc-tag, SV40NLS
1695..1724: Myc-tag
1731..1745: SV40NLS
2603..3191: Woodchuck hepatitis virus posttranscriptional regulatory element (WPRE)
3658..4246(complement): pUC ori
4446..5306(complement): beta lactamase

**Plasmid name: pEFs-Myc-bpNLS-Cas3d-bpNLS-6xHis**

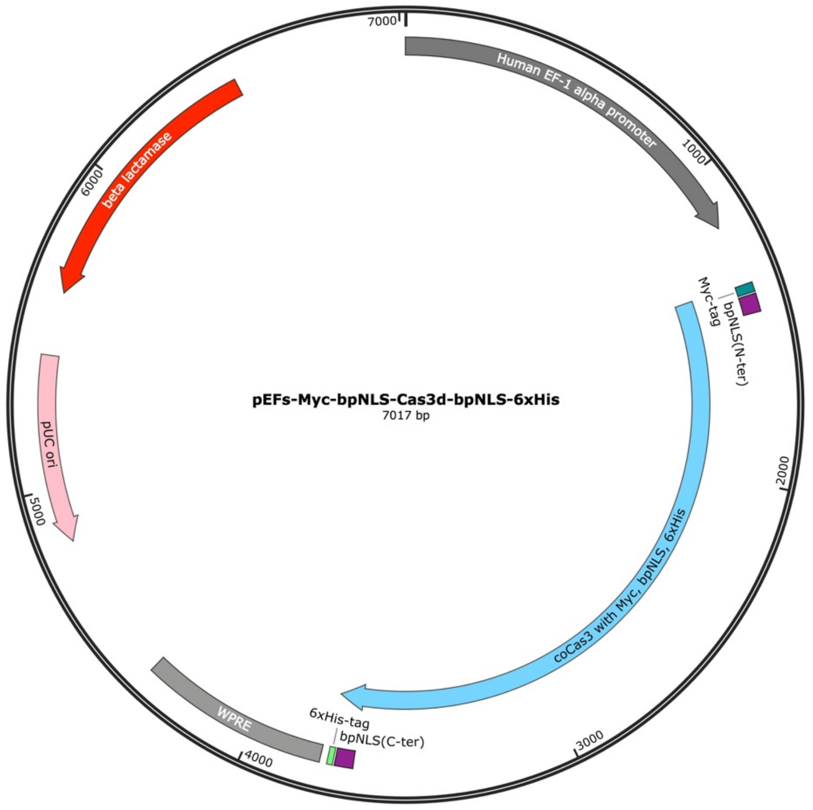


1..1182: Human EF-1 alpha promoter
1369..3753: coCas3 with Myc, bpNLS, 6xHis
1375..1404: Myc-tag
1411..1467: bpNLS(N-ter)
3670..3726: bpNLS(C-ter)
3733..3750: 6xHis-tag
3774..4362: Woodchuck hepatitis virus posttranscriptional regulatory element (WPRE)
4829..5417(complement): pUC ori
5617..6477(complement): beta lactamase

**Plasmid name: pEFs-Myc-bpNLS-Cas5d-bpNLS-6xHis**

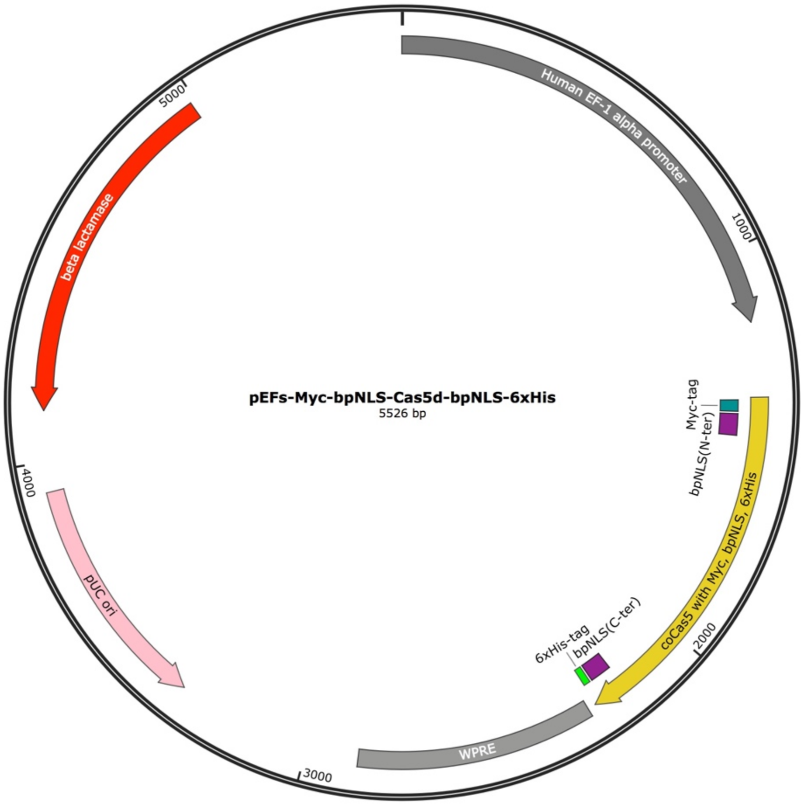


1..1182: Human EF-1 alpha promoter
1369..2262: coCas5 with Myc, bpNLS, 6xHis
1375..1404: Myc-tag
1411..1467: bpNLS(N-ter)
2179..2235: bpNLS(C-ter)
2242..2259: 6xHis-tag
2283..2871: Woodchuck hepatitis virus posttranscriptional regulatory element (WPRE)
3338..3926(complement): pUC ori
4126..4986(complement): beta lactamase

**Plasmid name: pEFs-Myc-bpNLS-Cas6d-bpNLS-6xHis**

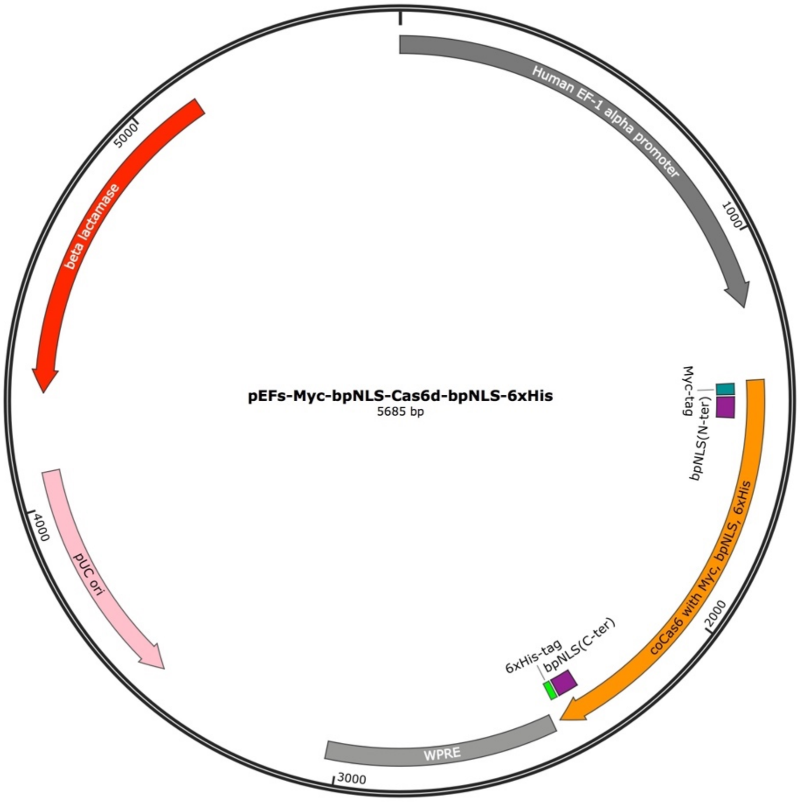


1..1182: Human EF-1 alpha promoter
1369..2421: coCas6 with Myc, bpNLS, 6xHis
1375..1404: Myc-tag
1411..1467: bpNLS(N-ter)
2338..2394: bpNLS(C-ter)
2401..2418: 6xHis-tag
2442..3030: Woodchuck hepatitis virus posttranscriptional regulatory element (WPRE)
3497..4085(complement): pUC ori
4285..5145(complement): beta lactamase

**Plasmid name: pEFs-Myc-bpNLS-Cas7d-bpNLS-6xHis**

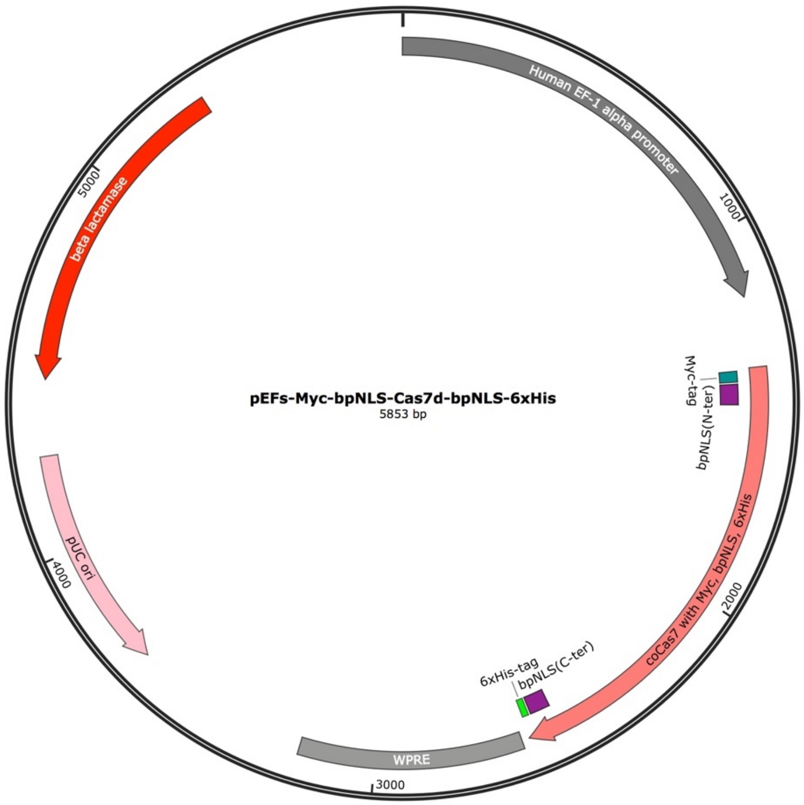


1..1182: Human EF-1 alpha promoter
1369..2589: coCas7 with Myc, bpNLS, 6xHis
1375..1404: Myc-tag
1411..1467: bpNLS(N-ter)
2506..2562: bpNLS(C-ter)
2569..2586: 6xHis-tag
2610..3198: Woodchuck hepatitis virus posttranscriptional regulatory element (WPRE)
3665..4253(complement): pUC ori
4453..5313(complement): beta lactamase

**Plasmid name: pEFs-Myc-bpNLS-Cas10d-bpNLS-6xHis**

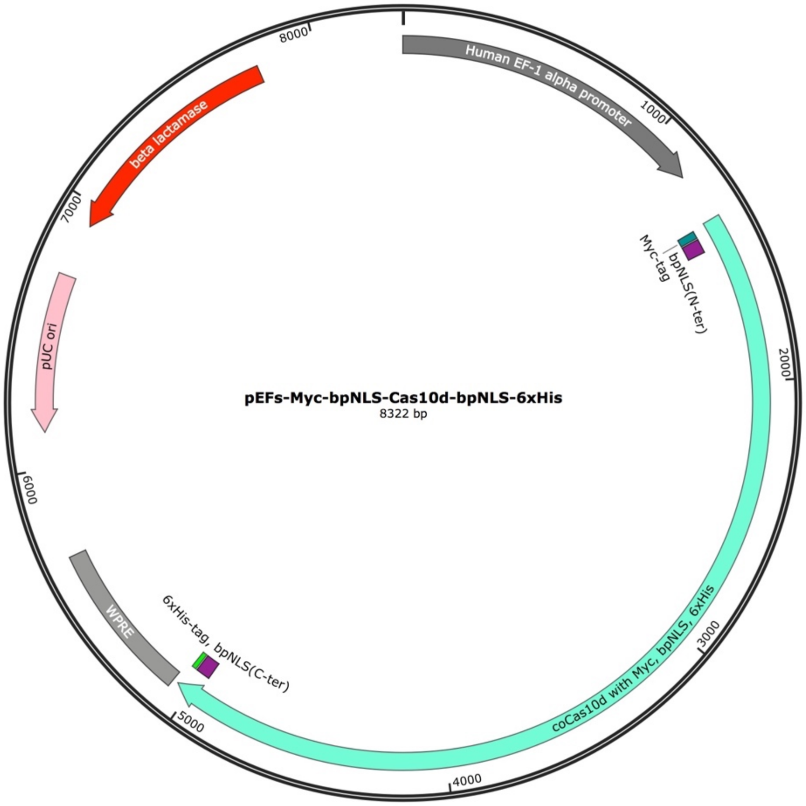


1..1182: Human EF-1 alpha promoter
1369..5058: coCas10d with Myc, bpNLS, 6xHis
1375..1404: Myc-tag
1411..1467: bpNLS(N-ter)
4975..5031: bpNLS(C-ter)
5038..5055: 6xHis-tag
5079..5667: Woodchuck hepatitis virus posttranscriptional regulatory element (WPRE)
6134..6722(complement): pUC ori
6922..7782(complement): beta lactamase

**Plasmid name: pEFs-Myc-bpNLS-Cas10d(H177A)-bpNLS-6xHis**

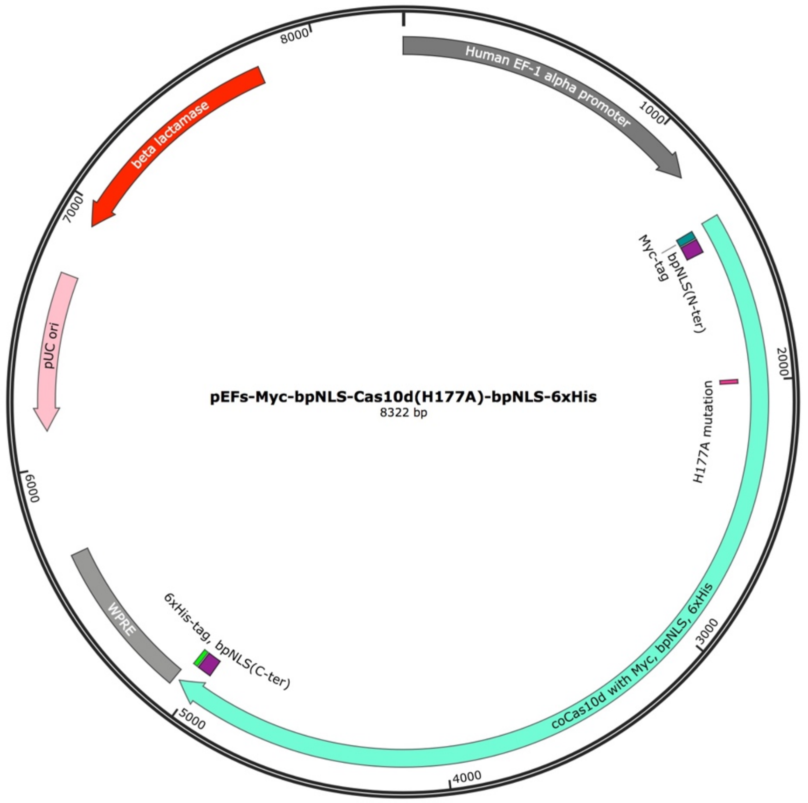


1..1182: Human EF-1 alpha promoter
1369..5058: coCas10d(H177A) with Myc, bpNLS, 6xHis
1375..1404: Myc-tag
1411..1467: bpNLS(N-ter)
1999..2001: H177A mutation
4975..5031: bpNLS(C-ter)
5038..5055: 6xHis-tag
5079..5667: Woodchuck hepatitis virus posttranscriptional regulatory element (WPRE)
6134..6722(complement): pUC ori
6922..7782(complement): beta lactamase

**Plasmid name: pEFs-6xHis-Myc-bpNLS-Cas7d-bpNLS**

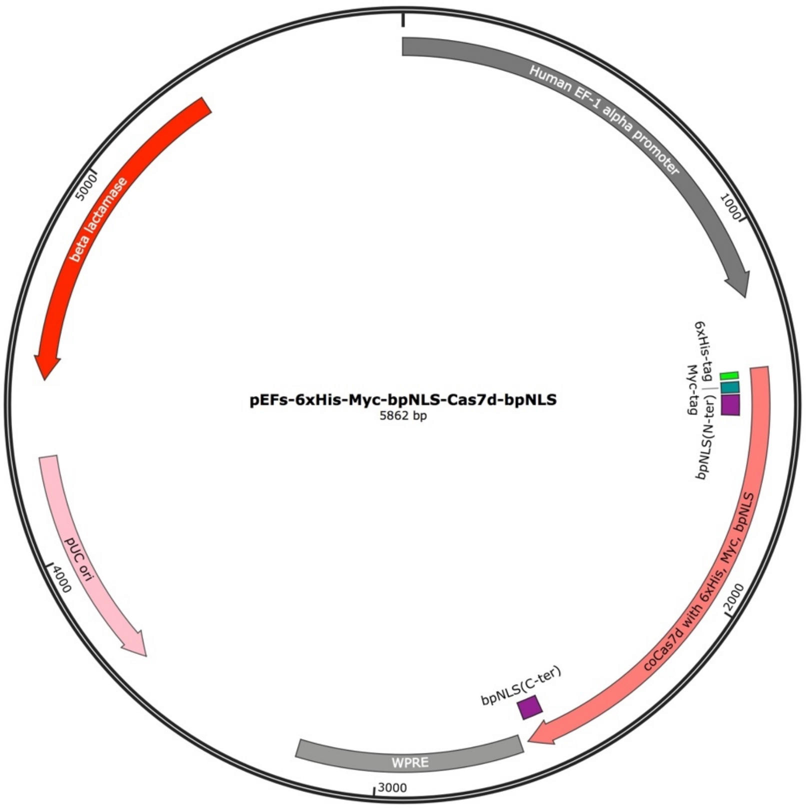


1..1182: Human EF-1 alpha promoter
1369..2598: coCas7d with 6xHis, Myc, bpNLS
1375..1392: 6xHis-tag
1402..1431: Myc-tag
1438..1494: bpNLS(N-ter)
2533..2589: bpNLS(C-ter)
2619..3207: Woodchuck hepatitis virus posttranscriptional regulatory element (WPRE)
3674..4262(complement): pUC ori
4462..5322(complement): beta lactamase

**Plasmid name: pAEX-hU6crRNA**

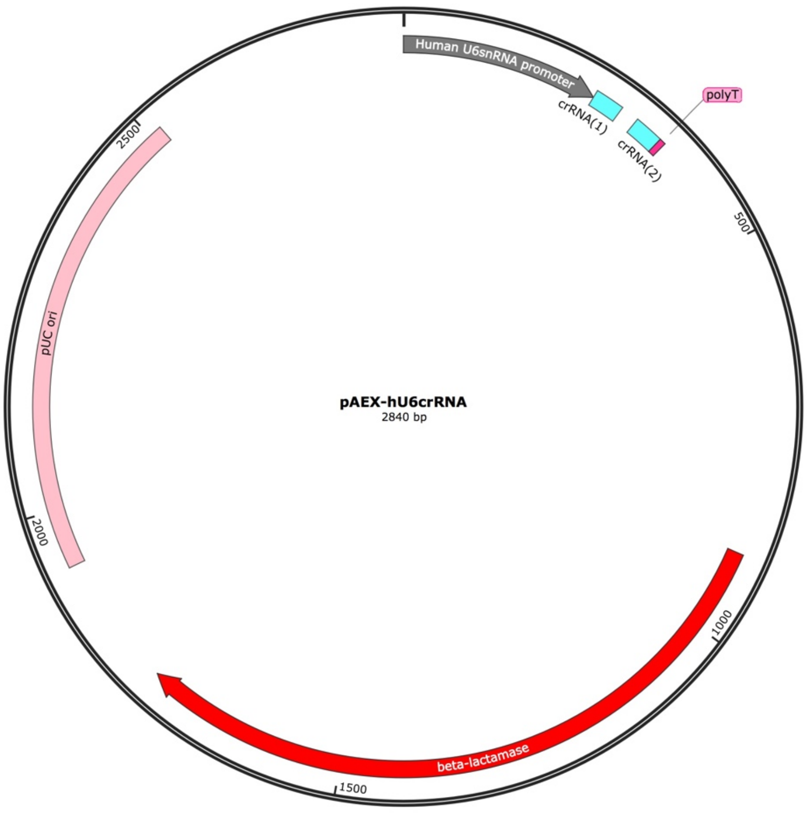


1..249: Human U6 snRNA promoter
250..286: crRNA(1)
310..346: crRNA(2)
347..354: polyT
897..1757: beta-lactamase
1928..2516: pUC ori

**Plasmid name: pAEX-hU6crRNA_mature**

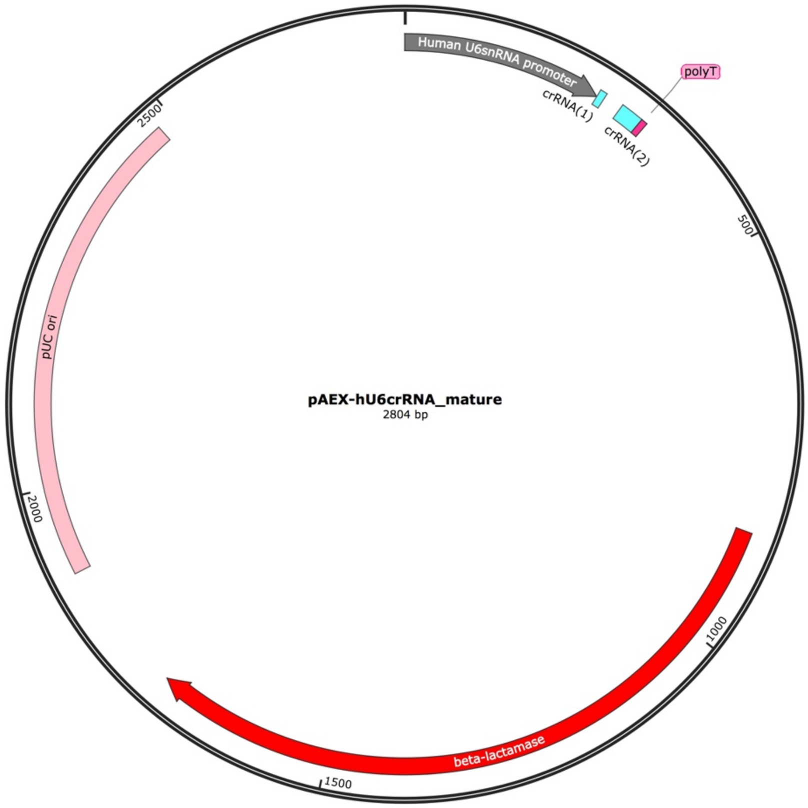


1..249: Human U6 snRNA promoter
250..258: Processed-crRNA(1)
282..310: Processed-crRNA(2)
311..318: polyT
861..1721: beta-lactamase
1892..2480: pUC ori

**Plasmid name: pCAG-nLUxxUC**

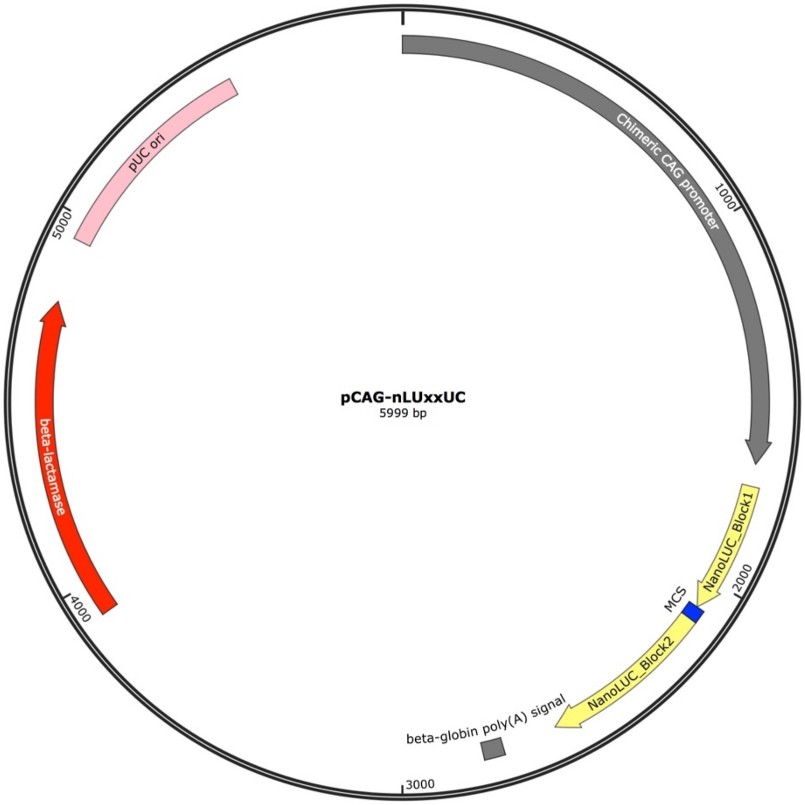


1..1665: Chimeric CAG promoter
1726..2079: NanoLUC_Block1 (#1 to #351 of NanoLUC with synthetic stop codon)
2080..2115: MCS
2116..2580: NanoLUC_Block2 (#52 to the original stop codon of NanoLUC)
2729..2784: rabbit beta-globin poly(A) signal
3912..4772: beta lactamase
4943..5531: pUC ori

**Plasmid name: pCAG-nLUxxUC-Block1-MCS**

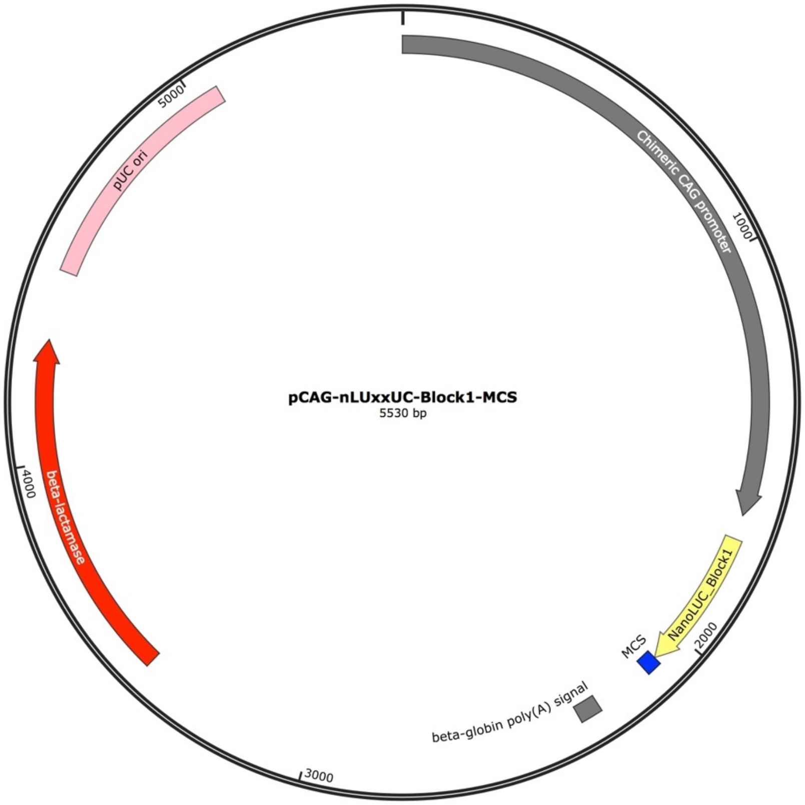


1..1665: Chimeric CAG promoter
1726..2079: NanoLUC_Block1 (#1 to #351 of NanoLUC with synthetic stop codon)
2080..2117: MCS
2260..2315: rabbit beta-globin poly(A) signal
3443..4303: beta lactamase
4474..5062: pUC ori

**Plasmid name: pCAG-nLUxxUC-MCS-Block2**

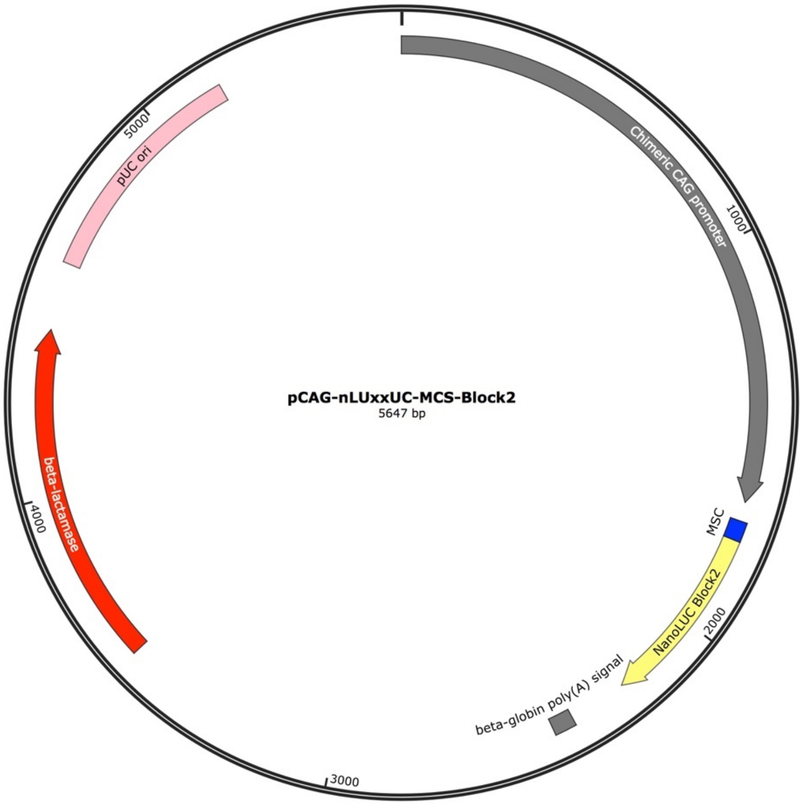


1..1665: Chimeric CAG promoter
1714..1763: MCS
1764..2228: NanoLUC_Block2 (#52 to the original stop codon of NanoLUC)
2377..2432: rabbit beta-globin poly(A) signal
3560..4420: beta lactamase
4591..5179: pUC ori

Supplemental Table S1. The list of TiD gRNAs target sequences.

| gRNA name | PAM | Sequence, 5'-3' | Target gene | Locus | Description |
|---|---|---|---|---|---|
| AAVS1_GTC_58-95(+) | GTC | accaatcctgtccctagtggccccactgtggggtg | *Hs* AAVS1 | Chr19: 55115748-55115782 | "GTC58-95" was used in Fig. 2. |
| AAVS1_GTC_70-107(+) | GTC | cctagtggccccactgtggggtggaggggacagat | *Hs* AAVS1 | Chr19: 55115760-55115794 | "AAVS70-107" was used in Fig. 1 and Fig. 2, and "GTC70-107" in Fig. 2. |
| AAVS1_GTC_159-196(+) | GTC | ccagctcggggacacaggatccctggaggcagcaa | *Hs* AAVS1 | Chr19: 55115849-55115883 | "AAVS159-196" and "GTC159-196" was used in Fig. 2. |
| AAVS1_GTC_180-217(-) | GTC | cacttcaggacagcatgtttgctgcctccagggat | *Hs* AAVS1 | Chr19: 55115901-55115867 | "GTC180-217" was used in Fig. 2. |
| EMX1_GTT_3(+) | GTT | CCAGAACCGGAGGACAAAGTACAAACGGCAGAAGCT | *Hs* EMX1 | Chr2: 72918572-72918607 | "GTT_3" was used in Fig. 2 and "GTT_3(+)" in Fig. 3. |
| EMX1_GTT_8(-) | GTT | TGTACTTTGTCCTCCGGTTCTGGAACCACACCTTCA | *Hs* EMX1 | Chr2: 72933823-72933788 | "GTT_8" was used in Fig. 2. |
| EMX1_GTT_9(-) | GTT | GATGTGATGGGAGCCCTTCTTCTTCTGCTCGGACTC | *Hs* EMX1 | Chr2: 72933888-72933853 | "GTT_9" was used in Fig. 2 and Fig. 3, and "GTT_9(-)" in Fig. 3. |
| EMX1_GTC_2(+) | GTC | CGAGCAGAAGAAGAAGGGCTCCCATCACATCAACCG | *Hs* EMX1 | Chr2: 72933858-72933893 | "GTC_2" was used in Fig. 2. |
| EMX1_GTC_9(-) | GTC | CTCCCCATTGGCCTGCTTCGTGGCAATGCGCCACCG | *Hs* EMX1 | Chr2: 72933927-72933892 | "GTC_9" was used in Fig. 2. |
| EMX1_GTC_10(-) | GTC | ATTGGAGGTGACATCGATGTCCTCCCCATTGGCCTG | *Hs* EMX1 | Chr2: 72933948-72933913 | "GTC_10" was used in Fig. 2. |
| EMX1_GTA_1(+) | GTA | CAAACGGCAGAAGCTGGAGGAGGAAGGGCCTGAGTC | *Hs* EMX1 | Chr2: 72918367-72918402 | "GTA_1" was used in Fig. 2. |
| EMX1_GTA_7(-) | GTA | CTTTGTCCTCCGGTTCTGGAACCACACCTTCACCTG | *Hs* EMX1 | Chr2: 72933819-72933784 | "GTA_7" was used in Fig. 2. |

Supplemental Table S2. The list of target DNA fragments used in the nanoLuc assay.

| | Sequence, 5'-3' | Locus |
|---|---|---|
| EMX1_GTT_3 | CCAGAACCGGAGGACAAAGTACAAACGGCAGAAGCT | Chr2: 72918572-72918607 |
| EMX1_GTT_8 | TGTACTTTGTCCTCCGGTTCTGGAACCACACCTTCA | Chr2: 72933823-72933788 |
| EMX1_GTT_9 | GATGTGATGGGAGCCCTTCTTCTTCTGCTCGGACTC | Chr2: 72933888-72933853 |
| EMX1_GTC_2 | CGAGCAGAAGAAGAAGGGCTCCCATCACATCAACCG | Chr2: 72933858-72933893 |
| EMX1_GTC_9 | CTCCCCATTGGCCTGCTTCGTGGCAATGCGCCACCG | Chr2: 72933927-72933892 |
| EMX1_GTC_10 | ATTGGAGGTGACATCGATGTCCTCCCCATTGGCCTG | Chr2: 72933948-72933913 |
| EMX1_GTA_1 | CAAACGGCAGAAGCTGGAGGAGGAAGGGCCTGAGTC | Chr2: 72918367-72918402 |
| EMX1_GTA_7 | CTTTGTCCTCCGGTTCTGGAACCACACCTTCACCTG | Chr2: 72933819-72933784 |
| AAVS1_58-95(+) | ACCAATCCTGTCCCTAGTGGCCCCACTGTGGGGTG | Chr19: 55115748-55115782 |
| AAVS1_70-107(+) | CCTAGTGGCCCCACTGTGGGGTGGAGGGGACAGAT | Chr19: 55115760-55115794 |
| AAVS1_159-196(+) | CCAGCTCGGGGACACAGGATCCCTGGAGGCAGCAA | Chr19: 55115849-55115883 |
| AAVS1_180-217(-) | CACTTCAGGACAGCATGTTTGCTGCCTCCAGGGAT | Chr19: 55115901-55115867 |

Supplemental Table S3. Primers for short-range PCRs to detect small in/dels

| Oligor name | Sequence, 5'-3' | Purpose | Description |
|---|---|---|---|
| EMX1_HMA1F | AGCCTCAGTCTTCCCATCAGGCTCT | HMA and cloning of the target region for *hEMX1* | PCR primers were used in Fig. 3. |
| EMX1-HMA1R | CCATGACTCCAGGCCTCCCCAAAG | HMA and cloning of the target region for *hEMX1* | |

Supplemental Table S4. Primers for long-range PCRs to detect large deletions

| Oligor name | Sequence, 5'-3' | Purpose | Description |
|---|---|---|---|
| hEMX1_19995_FW | CAACAGCTAATTCTGTCAAAATAATCCATC | Long PCR, Fig. 3a lane 1 (1st PCR ) | All data in Fig. 3a, b, and c, and Supplemental Fig S5 were detected by the nested PCR. |
| hEMX1_19995_RV | TCATAAGGTCTTATTCTTGTGCACTTTATC | Long PCR, Fig. 3a lane 1 (1st PCR ) | |
| hEMX1_17962_FW | TATTTATACACATGTTTCTTTAGCAAGGGA | Long PCR, Fig. 3a lane 2 (1st PCR ), lane 1 (nested PCR ) | |
| hEMX1_17962_RV | AGACATTTATTGACACCTACTCTAACTGAG | Long PCR, Fig. 3a lane 2 (1st PCR ), lane 1 (nested PCR ) | |
| hEMX1_14915_FW | ATTAACTATCAAAACTCAGGGCTAATCTTC | Long PCR, Fig. 3a lane 3 (1st PCR ), lane 2 (nested PCR ) | |
| hEMX1_14915_RV | CTTATCTATAAAATGCCACGAACAGCATCA | Long PCR, Fig. 3a lane 3 (1st PCR ), lane 2 (nested PCR ) | |
| hEMX1_12397_FW | CCCATACAAATACACACTAACTTTATCAGT | Long PCR, Fig. 3a lane 4 (1st PCR ), lane 3 (nested PCR ) | |
| hEMX1_12397_RV | CAGCTCCATGTCATTAGAGAATAGAGAG | Long PCR, Fig. 3a lane 4 (1st PCR ), lane 3 (nested PCR ) | |
| hEMX1_9109_FW | GTCAGATGATAGCATAGGTACACATTAGAT | Long PCR, Fig. 3a lane 5 (1st PCR ), lane 4 (nested PCR ) | |
| hEMX1_9109_RV | GTAATTGGTTAAACCTGTTCCGATGTCTG | Long PCR, Fig. 3a lane 5 (1st PCR ), lane 4 (nested PCR ) | |
| hEMX1_5385_FW | GTGTGTAGATTTTGTTTTTCTATGGTTGTG | Long PCR, Fig. 3a lane 5 (nested | |
| hEMX1_5385_RV | TTAATCTGGACTTATCCATGTTAGGACTT | Long PCR, Fig. 3a lane 5 (nested | |
| AAVS_24263_FW | GGCTTCAAATGTTTCAAAAACACATCAT | Long PCR, Fig. 3b lane 1 (1st PCR ) | |
| AAVS_24263_RV | TAAAAAGTGACACGTAAAGTTTCGTATGG | Long PCR, Fig. 3b lane 1 (1st PCR) | |
| AAVS_19031_FW | TTAGGAGATATACCTAATGTAAATGACGAG | Long PCR, Fig. 3b lane 2 (1st PCR), lane 1 (nested PCR ) | |
| AAVS_19031_RV | AAAGTTATGAGAACTGTAGAGAGTGAGTTG | Long PCR, Fig. 3b lane 2 (1st PCR), lane 1 (nested PCR ) | |
| AAVS_16624_FW | CTTAGCATAATGTCCTCAAGATACATCTAC | Long PCR, Fig. 3b lane 3, 4, 5 (1st PCR), lane 2 (nested PCR ) | |
| AAVS_16624_RV | GGATAACTAAAAAGAAGTGCATAAAAGAGT | Long PCR, Fig. 3b lane 3 (1st PCR), lane 2 (nested PCR ) | |
| AAVS_14147_RV | GATATGTAACCATTATTCTAGATGGCTATG | Long PCR, Fig. 3b lane 4 (1st PCR), lane 3 (nested PCR ) | |
| AAVS_11970_RV | ATAAACACAAACTCATAAACAACATACATC | Long PCR, Fig. 3b lane 5, 6, 7, 8, 9 (1st PCR), lane 4 (nested PCR ) | |
| AAVS_3585-3612_FW | GGGTCCAAGGGAAAAGGAGGACTGATCC | Long PCR, Fig. 3b lane 6, 7, 8, 9 (1st PCR), lane 3, 4, 5 (nested PCR ) | |
| AAVS_8359FW | CACAAATCTATCAAAAAGTTAAAAGCTG | Long PCR, Fig. 3b lane 6 (nested | |
| AAVS_8359RV | AAACAAAACTACTGACAAGTTGCTCTAC | Long PCR, Fig. 3b lane 5, 6 (nested | |
| AAVS_7077FW | ATTTCCTCGTCAGTCTCCCTTTCCTTCC | Long PCR, Fig. 3b lane 7 (nested | |
| AAVS_7077RV | TAGGGAGTGGAGTGTGGATTTCTCTTGC | Long PCR, Fig. 3b lane 7 (nested | |
| AAVS_5000FW | CTCCTAGATCCACGGGATAAATTAC | Long PCR, Fig. 3b lane 8 (nested | |
| AAVS_5000RV | GCTTATTTCTAGTTAAGGGGTCAGG | Long PCR, Fig. 3b lane 8 (nested | |
| AAVS_3594FW | CCCTCCGGCCTGTAGACTCCATTTC | Long PCR, Fig. 3b lane 9 (nested | |
| AAVS_3594RV | GCTCTGTTCAGCCCTAAGAATCCTG | Long PCR, Fig. 3b lane 9 (nested | |
